# ATAC and SAGA histone acetyltransferase modules facilitate transcription factor binding to nucleosomes in an acetylation independent manner

**DOI:** 10.1101/2023.10.27.564358

**Authors:** Kristin Chesnutt, Gizem Yayli, Christine Toelzer, Khan Cox, Gunjan Gautam, Imre Berger, László Tora, Michael G. Poirier

## Abstract

Transcription initiation involves the coordination of multiple events, starting with activators binding specific DNA target sequences, which recruits transcription co-activators to open chromatin and enable binding of general transcription factors and RNA polymerase II to promoters. Two key human transcriptional coactivator complexes, ATAC (ADA-Two-A-Containing) and SAGA (Spt-Ada-Gcn5-acetyltransferase), target genomic loci to increase promoter accessibility. To better understand the function of ATAC and SAGA histone acetyltransferase (HAT) complexes, we used *in vitro* biochemical and biophysical assays to characterize human ATAC and SAGA HAT module interactions with nucleosomes and how a transcription factor (TF) coordinates these interactions. We found that ATAC and SAGA HAT modules bind nucleosomes with high affinity, independent of post-translational modifications (PTMs) and TFs. ATAC and SAGA HAT modules directly interact with the VP16 activator domain and a TF containing this domain enhances HAT module acetylation activity. Surprisingly, ATAC and SAGA HAT modules increase TF binding to its DNA target site within the nucleosome by an order of magnitude independent of histone acetylation. Altogether, our results reveal synergistic coordination between HAT modules and a TF, where ATAC and SAGA HAT modules: (i) acetylate histones to open chromatin, and (ii) facilitate TF targeting within nucleosomes independently of their acetylation activity.

## INTRODUCTION

The eukaryotic genome is packaged in the nucleus as a highly condensed structure called chromatin. The basic repeating unit of chromatin is the nucleosome, which contains ∼147 base pairs (bp) of DNA wrapped ∼1.65 times around two copies of each histone protein (H2A, H2B, H3, and H4) (1–3). Human genomic DNA is wrapped into ∼20 million nucleosomes, which compact into dynamic higher-order structures that regulate DNA accessibility and controls genome functions, including transcription and DNA repair/replication (4, 5). Access to DNA is critical for DNA-mediated events (6). For example, transcription factors (TFs) must access specific DNA target sequences within promoters and enhancers to initiate the activation of genes by facilitating the recruitment of co-activators, including chromatin-modifying complexes (7, 8). However, the mechanisms by which TFs and chromatin-modifying complexes coordinate their gene-activating functions are not yet fully understood.

Chromatin-modifying complexes recognize (readers), deposit (writers) and remove (erasers) post-translational modifications (PTMs) of histones in a dynamic manner. Two highly conserved large multi-subunit transcriptional regulatory co-activator complexes that can acetylate histones at distinct residues are ATAC (ADA-Two-A-Containing) and SAGA (Spt-Ada-Gcn5-acetyltransferase) (9).

The metazoan ATAC and SAGA coactivator complexes contain 10 and 18-20 well-characterized subunits, respectively, which are organized in functional modules (10–12). SAGA is conserved from yeast to humans and organized into the histone acetyltransferase (HAT), histone H2Bub1 deubiquitinase (DUB), activator binding (AM), splicing (SM) and core modules (10–16). ATAC, which has been described in metazoans but is not found in yeast, also contains a HAT module (17) and a core module (18); however, the structural organization of ATAC is not yet known.

Importantly, ATAC and SAGA HAT modules (hereafter ATAC_HAT_ and SAGA_HAT_) are highly similar in that three of their subunits KAT2A/KAT2B, TADA3 and SGF29 are shared between them. The fourth and distinctive subunit is either TADA2A, or TADA2B for ATAC, or SAGA HAT modules, respectively (**Figure 1A**) (12, 19, 20). The similar HAT modules of these two endogenous co-activator complexes seem to have different histone acetylation targets *in vivo*, where SAGA acetylates H3K9 and H3K14 (17, 21, 22), while it has been suggested that ATAC acetylates both histone H3 and H4 (16, 23–27). In addition, ATAC and SAGA play different regulatory roles in transcription regulation and/or cellular homeostasis (25, 27–34). How these two coactivator complexes containing very similar HAT modules target histone acetylation differentially remains an open question.

**Figure 1.**
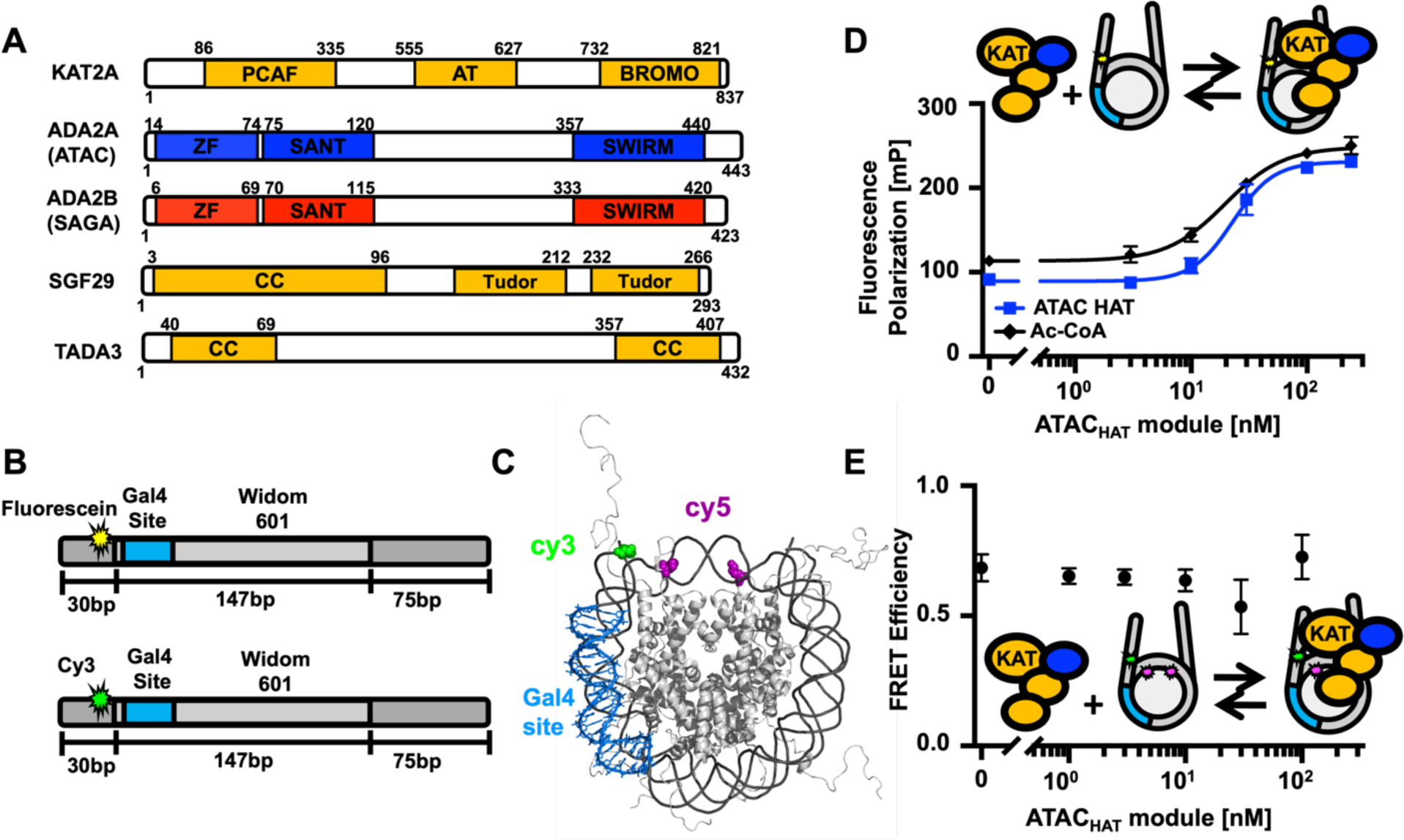
ATAC and SAGA HAT module organization and ATAC HAT module interactions with nucleosomes. (A) Schematic representations of the shared subunits of the ATAC and SAGA HAT modules (KAT2A, SGF29 and TADA3), together with ADA2A and ADA2B, which are ATAC and SAGA subunits, respectively (10). PCAF: PCAF homology domain; AT: Acetyl-transferase domain; BROMO: bromo domain; ZF: zinc-finger domain; SANT: <u>S</u>wi3, <u>A</u>DA2, <u>N</u>-Cor and <u>T</u>FIIIB domain; Tudor: Tudor domain; CC: coiled coil domain. (B) (Upper part) Schematic of the DNA used in Fluorescence Polarization experiments. DNA is 252 bp long and consists of the Widom 601 sequence (light gray) flanked by a 30 bp (left, dark gray) and a 75 bp linker DNA (right, dark gray). Within the 601 sequence is a 19 bp Gal4 binding site (light blue) at 8-26 bp in the 601 nucleosome positioning sequence (NPS). Fluorescein (yellow) is attached to a modified thymine 4 bp into the 30 bp linker DNA away from the 601 NPS. (Lower part) Schematic of the DNA used in FRET experiments. Same sequence as above but with Cy3 (green) attached to a modified thymine 4 bp into the 30 bp linker DNA away from the 601 NPS. (C) Nucleosome structure (PDB 1KX5) indicating the estimated difference between Cy3 on the DNA (green) and Cy5 on H2A (magenta). The histone octamer is gray, DNA is dark gray, and Gal4 binding site is blue. (D) Fluorescence Polarization in mP of nucleosomes with increasing amounts of the ATAC_HAT_ module. The ATAC_HAT_ module binds with an S_1/2_ of 23 ± 4 nM (blue square), reaching saturating conditions around 100 nM. The ATAC_HAT_ module with Ac-CoA binds at a S_1/2_ of 19.6 ± 0.6 nM (black diamond). (E) FRET efficiency of the Cy3-Cy5 labeled nucleosome with increasing concentrations of ATAC_HAT_.

Recent cryo-EM studies (15, 16, 35) have revealed the structure of most of the SAGA complex at high resolution. However, SAGA_HAT_ could not be resolved in the structures, indicating that the module has a high degree of flexibility within the SAGA complex. This flexibility is further enhanced by nucleosome binding, which appears to result in the displacement of the HAT module (16). In addition, a metazoan SAGA_HAT_ has been reported to function on its own as the ADA complex (36) and with the addition of the protein Chiffon (37).

Targeting of the SAGA complex by TFs occurs at least in part through interactions with the AM module, which in humans is TRRAP (36, 38). In addition, Myc has been described to interact directly with TRRAP and KAT2A (39–42). Interestingly, the SAGA_HAT_ module has a number of nucleosome binding domains indicating that it can target nucleosomes directly (10). This in combination with the observation that the AM module is not included in the ATAC complex, nor in the ADA complex, raises questions about how TFs target these endogenous co-activator complexes to chromatin.

Uncovering how these human HAT modules engage with TFs to target, acetylate and open chromatin is important for understanding mechanisms by which these co-activator complexes function to facilitate transcription initiation. Here we report systematic biochemical and biophysical studies of the interactions of SAGA_HAT_ and ATAC_HAT_ with the activator Gal4-VP16 and nucleosomes that contain the Gal4 target sequence. We find that both HAT modules directly interact strongly with nucleosomes and the VP16 activator. We then asked how the chimeric TF, Gal4-VP16, functions with each HAT module to open and acetylate nucleosomes. We focused on the Gal4 target site located in the nucleosome entry-exit region, a common position for TF binding sites (43–45). We determined that Gal4-VP16 significantly enhances histone acetylation of both ATAC_HAT_ and SAGA_HAT_, with a larger impact on ATAC_HAT_, and that this depends on the VP16 activator domain (AD). We then investigated the impact of each HAT module on Gal4-VP16 binding to its target site within nucleosomes. Interestingly, we found that both ATAC_HAT_ and SAGA_HAT_ increase Gal4-VP16 binding by an order of magnitude, but these increases did not depend on the histone acetylation activity of either complex. These results indicate that both ATAC_HAT_ and SAGA_HAT_ function with TFs to not only acetylate nucleosomes, but they also facilitate TF binding within nucleosomes independent of their histone acetylation activity.

## MATERIAL AND METHODS

### DNA preparation and purification

DNA constructs for reconstituting nucleosomes for FRET and acetylation assay experiments were prepared using PCR with Cy3-labeled oligonucleotides and a plasmid containing the Widom 601 nucleosome positioning sequence (NPS) with the Gal4 binding site (5’-CCGGAGGGCTGCCCTCCGG-3’) at bases 8-26 (46–48). Oligonucleotides were labeled with Cy3 NHS ester (GE Healthcare) at an amine-modified internal thymine and HPLC purified on a 218TP C18 reverse-phase column (Grace/Vydac) as previously described (49). DNA constructs for reconstituting nucleosomes for Fluorescence Anisotropy were prepared using PCR with Fluorescein-labeled oligonucleotides and a plasmid containing the Widom 601 nucleosome positioning sequence (NPS). Following PCR amplification, DNA was purified using a MonoQ column (GE healthcare).

### Core histone expression, purification, and histone octamer preparation

The histone octamer was prepared as previously described (49). Human histones: H2A(K119C), H2B, H3, and H4 were purchase from Histone Source. Lyophilized histones were resuspended in unfolding buffer (20 mM Tris-HCL pH 7.5, 7 M guanidinium, and 10 mM DTT) at 5 mg/ml for 1 hr then spun to remove aggregates. To determine concentration, absorption at 276 nm was measured for each unfolded histone. Histone octamers were refolded by combining (H2A and H2B):(H3 and H4) together at a ratio of 1.2:1. Double dialysis was performed to form the histone octamer into refolding buffer (10 mM Tris-HCl pH 7.5, 1 mM EDTA, 2 M NaCl, and 5 mM BME). The octamer was removed from dialysis and labeled with Cy5 maleimide (GE healthcare) at H2A(K119C) as previously described (50, 51). Labeled octamer was purified over a Superdex 200 (GE Healthcare) gel filtration column to remove excess dimer, tetramer, and dye. Labeling efficiency was determined by Ultraviolet-visible (UV-Vis) absorption, and purity of each octamer was confirmed by SDS-PAGE (**Supplementary Figure S1)**.

### Nucleosome preparation

Nucleosomes were reconstituted and purified as previously described (49, 52). DNA and histone octamers were combined at a ratio of 1.25:1 in high salt refolding buffer (0.5x TE pH 8.0, 2 M NaCl, 0.5 mM EDTA and 1 mM benzamidine−HCl) and reconstituted using double dialysis with 4 L of reconstitution buffer at 4 °C for 5-6 h and then changed into a new 4 L bucket of buffer at 4 °C overnight (0.5x TE pH 8.0, 0.5 mM EDTA, 1 mM benzamidine−HCl). Dialyzed nucleosomes were purified in a 5-30% w/v sucrose gradient with a SW41 Ti (Beckman Coulter) rotor in an Optima L-90K ultracentrifuge (Beckman Coulter) spinning at 41,000 rpm for 22 h at 4 °C. Sucrose gradients were fractionated into 0.4 mL fractions and analyzed by native polyacrylamide gel electrophoresis. Fractions containing center positioned nucleosomes were concentrated and buffer exchanged into 0.5x TE pH 8 with a 30 kDa centrifugal filter. To verify purity, nucleosomes were analyzed by a native 5% acrylamide gel and 0.3x TBE (**Supplementary Figure S1**).

### Recombinant Gal4-DBD and Gal4-VP16 expression and purification

The Gal4 DNA binding domain from amino acid 1 to 147 (hereafter called Gal4-DBD) (48, 53) and Gal4 DNA binding domain fused to the VP16 activator domain (hereafter called Gal4-VP16) (54) were expressed in Rosetta™ BL21 (DE3) pLysS cells. Cells were grown in 2xYT and induced at OD_600_ of 0.5 with 1 mM IPTG and 100 mM zinc acetate. Cells were expressed for 3 h then spun at 4000 g for 15 minutes. The supernatant was removed, and cell pellets were frozen with liquid nitrogen.

Cells were thawed and resuspended in 30 mL of Gal4 buffer A (50 mM Tris-HCl pH 8, 200 mM NaCl, 10 mM imidazole, 10 mM BME, 20 μM zinc acetate, 1 mM DTT, and 1 mM PMSF) with leupeptin and pepstatin added at a final concentration of 20 *μ*g/ml. Resuspended cells were sonicated and spun at 23,000g for 15 minutes. Lysate containing the protein was collected. The lysate was added to a Ni-NTA column and washed with Gal4 buffer A. The protein was eluted with Gal4 buffer B (50 mM Tris-HCl pH 7.5, 200 mM NaCl, 200 mM imidazole, 20 μM zinc acetate, 1 mM DTT, 1 mM PMSF, and 0.2% TWEEN 20). Fractions were run on a 12% acrylamide SDS gel, pooled, and dialyzed into Gal4 buffer C with 200 mM NaCl (25 mM Tris-HCl pH 7.5, 200 mM NaCl, 20 μM zinc acetate, 1 mM DTT, and 1 mM PMSF). The protein was further purified after dialysis using cation exchange chromatography with a gradient from 200 mM to 600 mM NaCl using Gal4 buffer C with appropriate salt concentrations. Fractions containing the protein were pooled, concentrated, exchanged into Gal4 buffer D (10 mM HEPES pH 7.5, 200 mM NaCl, 10% glycerol, 20 μM zinc acetate, 1 mM DTT, and 1 mM PMSF), and then flash frozen for storage at -80 °C. The final yield was ∼5 mg/L of culture grown. Purity was confirmed by SDS-PAGE (**Supplementary Figure S2**).

### ATAC and SAGA HAT module expression and purification

Recombinant ATAC and SAGA HAT modules were produced using the MultiBac system (55) as previously described (17). In short, a construct containing human KAT2A with a N-terminal histidine tag followed by a tobacco etch virus protease cleavage site, human TADA2A or TADA2B, mouse TADA3 and human SGF29 were integrated in the EMBacY bacmid via TN7 transposition (56). The resulting recombinant baculoviruses were then used to produce the corresponding HAT modules by infecting SF21 cells. Cells were harvested, flash-frozen in liquid nitrogen and stored at -80°C. For the purification, cell pellets of the respective HAT module were thawed and resuspended in HAT TALON buffer A (50 mM Trizma® base, 400 mM NaCl, 10 mM Imidazole, pH 8) supplemented with cOmplete™ EDTA-free protease inhibitor tablets (Roche) and Benzonase® nuclease. Resuspended cells were lysed using sonication and spun down at 40,000 x g for 45 minutes. The supernatant was additionally filtered through a folding filter and then incubated with HisPur™ Cobalt resin or loaded on a prepacked HiFliQ Cobalt-NTA FPLC column with a peristaltic pump. The protein was eluted via a step gradient of 7%, 30%, 50% and 100% HAT TALON buffer B (50 mM Trizma® base, 400 mM NaCl, 250 mM Imidazole, pH 8). The elutions were applied on an acrylamide SDS gel and fractions containing the complex were pooled accordingly. TCEP was added to a final concentration of 0.25 mM and the imidazole content diluted with HAT TALON buffer A where appropriate, to facilitate removal of the His-Tag by tobacco etch virus (TEV) protease cleavage of the His-Tag overnight. Samples were concentrated and loaded onto a Superdex 200 increase size exclusion chromatography (SEC) column equilibrated with HAT SEC buffer (25 mM Trizma® base, 400 mM NaCl, 0.25 mM TCEP, pH 8). Fractions were evaluated by SDS-PAGE (**Supplementary Figure S3**), pooled and concentrated. Aliquots were stored frozen in liquid nitrogen.

### Förster Resonance Energy Transfer efficiency measurements

For nucleosome-TF binding measurements, nucleosomes and TF were incubated with either the SAGA or ATAC HAT module and Ac-CoA in T75 buffer (10 mM Tris-HCl pH 8, 75 mM NaCl, 10% glycerol, and 0.25% Tween-20) in a 20 *μ*l volume at room temperature for at least 10 minutes and then analyzed using Horiba Scientific Fluoromax 4. Nucleosomes were at 1 nM in all experiments. For nucleosome-HAT module binding measurements, nucleosomes and either the ATAC or SAGA HAT module were incubated in T75 buffer in a 20 *μ*l volume at room temperature for at least 10 minutes as above. Fluorescence spectra were measured with Fluoromax 4 by exciting at 510 and 610 nm and measuring emission from 530 to 750 nm and 630 to 750 nm for donor and acceptor excitations, respectively. FRET efficiency was then calculated using the ratio_A_ method (57). FRET efficiency values were normalized against the FRET efficiency value of nucleosomes in the absence of titrant. The titrations were carried out in triplicate. The average FRET efficiency and standard error was determined for each TF concentration. The S_1/2_ and its standard error was then determined from a weighted fit to a non-cooperative binding isotherm using Prism 9 software.

### Fluorescence polarization measurements of HAT modules binding to nucleosomes

Fluorescent polarization measurements were carried out by mixing increasing amounts of either ATAC_HAT_ or SAGA_HAT_ with 5 nM NCP_FA_ in 75 mM NaCl, 25 mM Tris-HCl pH 7.5, and 0.25% Tween20 in a 30 *μ*l reaction volume. 0.3 mM Ac-CoA was added to buffer for measurements taken with Ac-CoA. Samples were loaded and incubated in a Corning round bottom polystyrene plate. Polarization measurements were acquired using a Tecan infinite M1000Pro plate reader by exciting at 470 nm. Polarized emission was measured at 519 nm with 5 nm excitation and emission bandwidths. Fluorescence polarization was calculated from the emission polarized parallel and perpendicular to the polarized excitation light as previously described (58, 59). The S_1/2_ was determined by fitting the data to a non-cooperative binding isotherm. S_1/2_ values were averaged over three separate experiments with error calculated as the standard error between the runs.

### Expression, Purification of Glutathione S-Transferase Fusion Proteins and HAT module Binding Assay

The expression and binding of Glutathione S-Transferase (GST) fusion proteins to glutathione Sepharose, and HAT binding assay (**Supplementary Figure S4**) was previously described (60). Briefly, *Escherichia coli*, DH5α were transformed with either pGEX-2T-GST or pGEX-2T-GST-VP16 vectors. Overnight cultures were diluted 1:100 in LB medium with ampicillin (100 mg/ml) and incubated at 37°C with shaking. After 1 h incubation, isopropyl-b-D-thiogalactopyranoside (IPTG) was added (final concentration 0.1 M) and the cultures were further incubated with shaking for 4 h. Cultures were pelleted by centrifugation at 5,000 g for 5 minutes at 4°C and pellets were resuspended in 1/10 volume of NETN buffer (20 mM Tris pH 8.0,100 mM NaCl, 1 mM EDTA, 0.5 % NP40), bacteria lysed by mild sonication and centrifuged at 10,000 g for 5 minutes at 4°C. Supernatants were loaded on Glutathione Sepharose 4B resin to immobilize either GST, or GST-VP16 fusion protein and incubated for 1h at 4°C. Following several washes of GST-, or GST-VP16-bound beads with NETN buffer; crude SF9 insect whole cell protein extract with no infection (as a negative control), SF9 cell extract expressing either the four ATAC_HAT_, or SAGA_HAT_ module subunits, were loaded and incubated overnight at 4°C, then washed with five times NETN buffer containing either 75 mM or 150 mM NaCl. The resins were boiled in 1x Laemmli buffer and bound proteins resolved by SDS-polyacrylamide gel electrophoresis. Proteins were visualized by western blot analyses with the following antibodies (as indicated in the corresponding Figures): anti-GCN5 2GC2C11 mouse monoclonal antibody (mab) (61), anti-TADA3 #2678 rabbit polyclonal antibody (pAb) (25), anti-SGF29 #2461 pAb (25), anti-VP16 #5GV2 mAb (62).

### HAT assays and corresponding Western blot assays

Either the ATAC or SAGA HAT module were incubated with Gal4-VP16 or Gal4-DBD, along with 1 nM of nucleosomes in T75 buffer with 30 μM Ac-CoA for 30 minutes. Reaction was quenched with 6x SDS loading buffer and samples were incubated in 95°C for 5 minutes before loading onto a 16% SDS-PAGE gel. Gels were then transferred onto a membrane and H2A-Cy5 labeled nucleosomes were fluorescently imaged to control for loading. Membranes were incubated with anti-H3K9ac (abcam 4441) and anti-H4K5ac (abcam 51997) antibodies at 1:1000 concentrations at 4°C overnight. The membranes were then probed with horseradish peroxidase-conjugated secondary antibodies and developed with enhanced chemiluminescence western blotting substrate (Thermo Scientific). Blots were visualized with detection system. The protein bands were quantified using the densitometry analysis in ImageJ software (NIH, Washington, DC, USA). Statistical analysis was performed using GraphPad Prism software. Two-tailed, unpaired t-test with Welch’s correction was conducted using Prism 9 software. Error bars in figures represent standard error of the mean.

## RESULTS

### Human ATAC and SAGA HAT modules interact with unmodified nucleosomes independent of the acetyl-CoA cofactor

While ATAC and SAGA complexes are reported to interact with histone tails of nucleosomes (10), the mechanism of ATAC_HAT_ and SAGA_HAT_ interactions with nucleosomes remain elusive. To characterize human ATAC_HAT_ and SAGA_HAT_ interactions with nucleosomes, Fluorescence Anisotropy (FA) was performed using a fixed concentration of unmodified nucleosomes and increasing concentrations of either the ATAC_HAT_ or SAGA_HAT_. Recombinant nucleosomes were prepared with a Widom 601 nucleosome positioning sequence (NPS) with a Gal4 binding site that starts at the eighth bp and extends 26 bp into the nucleosome (**Figure 1B**). The Gal4 binding site is included for TF binding studies described below. On the side proximal to the Gal4 binding site, the nucleosome was labeled with a fluorescein fluorophore that is attached to a 30 bp DNA extension at the eighth bp outside of the nucleosome, while the side of the nucleosome that is distal to the Gal4 binding site contained a 75 bp DNA extension. As either the ATAC_HAT_ or SAGA_HAT_ was titrated, an increase in polarization of the fluorescein fluorophore was observed, indicative of binding (59). We report HAT module binding to nucleosomes as the S_1/2_, the concentration of the HAT module required for half of the nucleosomes to be bound. FA measurements were fit to a binding isotherm with the ATAC_HAT_ binding with an S_1/2_ of 23 ± 4 nM and reaching saturating conditions around 100 nM (**Figure 1D**, **Supplementary Table S1**), while the SAGA_HAT_ S_1/2_ was 25.0 ± 0.3 nM, reaching saturating conditions around 100 nM (**Supplementary Figure S5A**). To determine the impact of acetylation catalysis on to the binding of the HAT modules to the nucleosome, we next added acetyl-CoA (Ac-CoA,) which is required for the acetylation activity of ATAC and SAGA (63). Titrations with Ac-CoA revealed that acetylation does not significantly impact the interaction of ATAC_HAT_ and SAGA_HAT_ modules with nucleosomes (**Figure 1D, Supplementary Figure S5A**). Overall, these results indicate that ATAC_HAT_ and SAGA_HAT_ modules strongly bind unmodified nucleosomes independent of acetylation and TFs.

### Binding of ATAC and SAGA HAT modules to the nucleosome does not significantly alter the wrapping of DNA into the nucleosome

Histone H3 tails are primary targets for acetylation by ATAC and SAGA and extend away from the nucleosome, where the DNA exits. Therefore, we next investigated if the binding of ATAC_HAT_ or SAGA_HAT_ would change the amount of DNA wrapped into the nucleosome. To this end, we prepared nucleosomes labeled with both Cy3 and Cy5 fluorophores as we have done previously for other H3 tail binding domains (58, 64). The DNA was labeled with Cy3 so it is located at the same locations as fluorescein, while the Cy5 was attached to histone H2AK119C (**Figure 1B-C**). These labeling positions undergo efficient energy transfer when the nucleosome is fully wrapped, while the FRET efficiency reduces significantly as the nucleosome partially unwraps (65). We carried out titrations with ATAC_HAT_ (**Figure 1E**) and SAGA_HAT_ (**Supplementary Figure S5B**) over the concentration range they bind nucleosomes. We found that the FRET reduction is minimal over the concentration range that they bind nucleosomes revealing that ATAC_HAT_ and SAGA_HAT_ binding does not directly impact nucleosome unwrapping.

### KAT2A, ATAC and SAGA HAT modules efficiently bind to the VP16 activator domain

Activators are known to interact with coactivator complexes. In the yeast SAGA complex, VP16 AD has been shown to specifically interact with the Tra1 subunit (66). Less is known about possible activator interactions with the mammalian ATAC_HAT_ or SAGA_HAT_ modules, nevertheless it has been shown that the human SAGA complex can bind to VP16 AD or to Myc (42, 60). To investigate whether the ATAC_HAT_ or SAGA_HAT_ modules directly interact with the VP16 AD, we tested their binding abilities to the activation domain of VP16 when fused to GST (**Figure 2A**). The GST-VP16 fusion protein was immobilized on glutathione-agarose resin in parallel with GST, as a negative control. ATAC_HAT_ and SAGA_HAT_ were able to bind to the GST-VP16 fusion protein, but not, or only very weakly, to GST alone (**Figure 2B-C**). Moreover, we examined whether KAT2A alone could bind the VP16 AD. Similar to the results obtained with purified HAT modules, recombinantly purified KAT2A is able to bind directly to the VP16 AD (**Supplementary Figure S4).** These results indicate that the HAT modules of ATAC and SAGA can specifically interact with the VP16 AD and this interaction is possibly through a direct interaction between VP16 AD and KAT2A, which is the shared enzymatic subunit of ATAC and SAGA HAT complexes.

**Figure 2.**
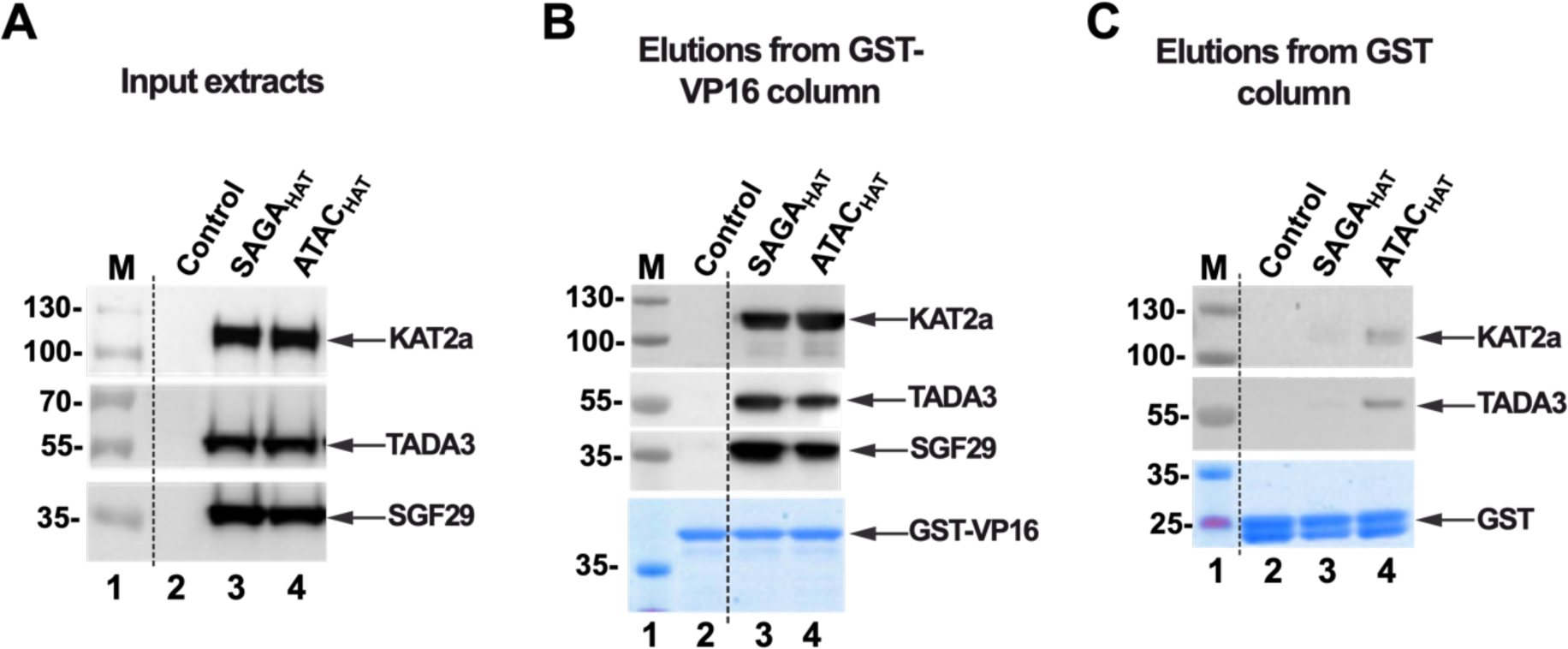
HAT module of SAGA and ATAC complexes directly interact with VP16 AD. (A) Image of immunoblot input extracts containing non-infected SF9 cell extract (Control, lane 2), and SF9 cell extracts expressing the subunits of either ATAC_HAT_, or SAGA_HAT_ (lane 3 and 4). Input extracts were incubated with either GST-VP16 (B) or GST (C) columns, washed and eluted. Elutions were resolved by SDS-PAGE and analyzed by Western blot with the indicated antibodies. Equal loading was tested by Coomassie brilliant blue staining of either GST-VP16 (B) or GST (C) protein containing beads.

### Gal4-VP16 binds its target site within nucleosomes via the nucleosome site-exposure model

Having determined that ATAC_HAT_ and SAGA_HAT_ can efficiently bind the VP16 AD, we proceeded to investigate the binding of the well-established model transcription factor Gal4-VP16 within nucleosomes alone (54). The Gal4-VP16 fusion TF contains the first 147 amino acids of the Gal4 transcription factor (Gal4-DBD), which contains the DNA binding and dimerization domains, and is fused to the highly potent transcription VP-16 AD (**Figure 3A**) (54). These measurements provide critical information for later measurements that include both Gal4-VP16 and either ATAC_HAT_ or SAGA_HAT_.

**Figure 3.**
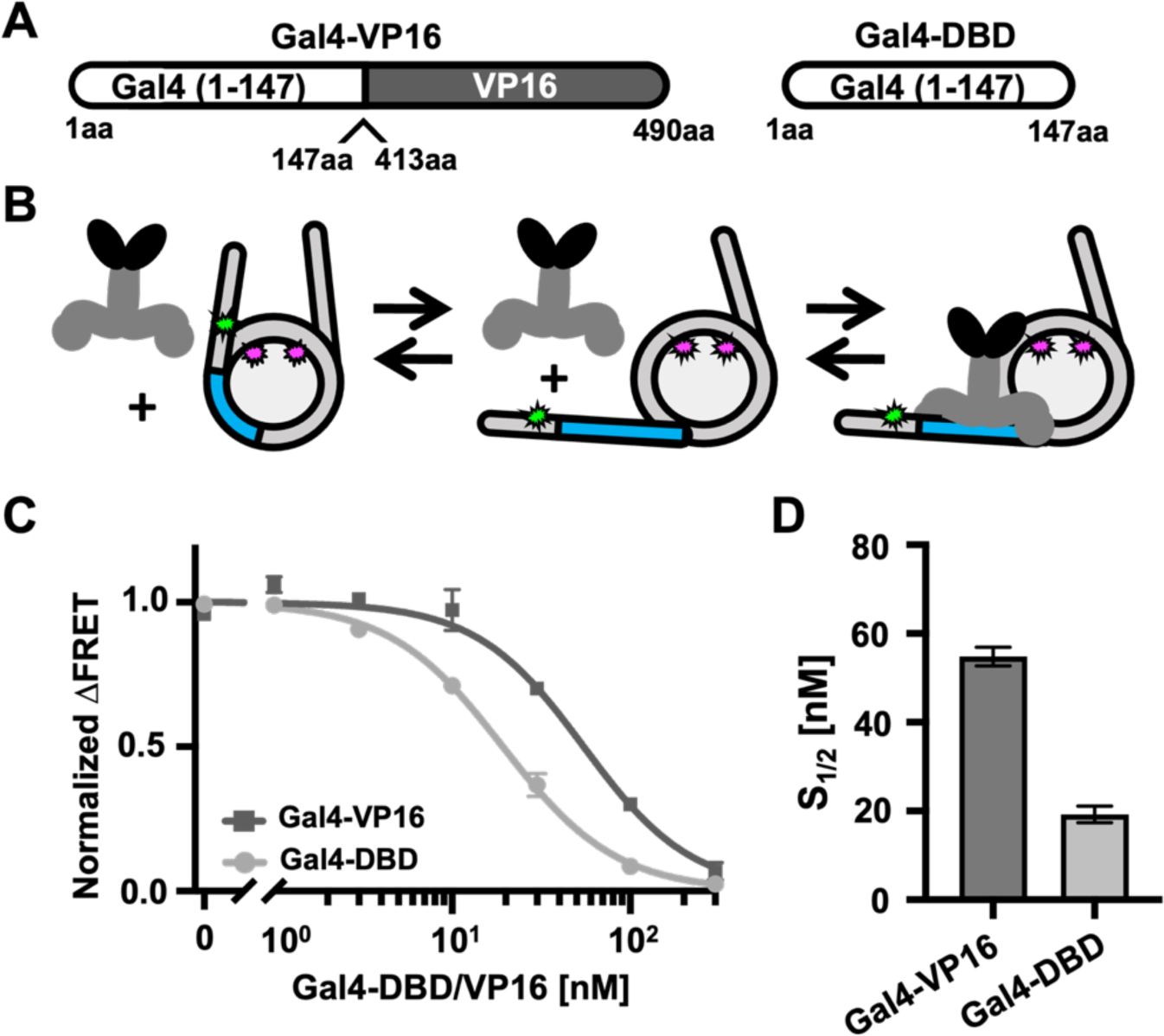
TFs interactions with nucleosomes. (A) Schematic of TFs: Gal4-VP16 and Gal4-DBD. Gal4-VP16 contains the DNA binding domain and dimerization domain of Gal4 fused to the VP16 activator domain from the Herpes Simplex Virus. Gal4-DBD contains the DNA binding domain and dimerization domain of Gal4. (B) Schematic of Gal4-VP16 binding equilibrium to a partially unwrapped nucleosome. (C) Normalized ΔFRET (the asymptote of the beginning of the hill plot fit set to 1 and the asymptote of the ending of the fit is set to 0) of Gal4-VP16 (dark gray squares) and Gal4-DBD (light gray circles) binding nucleosomes at increasing amounts. Gal4-VP16 binds with an S_1/2_ of 55 ± 2 nM and Gal4-DBD binds with an S_1/2_ of 19 ± 1 nM. (D) Bar plot of S_1/2_ and standard error values from the weighted fits to binding isotherms in C.

To detect Gal4-VP16 binding to nucleosomes, we used the Cy3-Cy5 labeled nucleosomes that were used to detect if the HAT modules directly induce nucleosome unwrapping (**Figure 1A-B**), which has been previously used to detect TF binding within nucleosomes (46, 50, 67). As mentioned above, the nucleosomes contained a 19 bp Gal4 binding site that starts at the eighth base pair and extends 26 base pairs into the Widom 601 NPS (**Figure 1B**). This positions the Gal4 binding site within the nucleosome near, where the DNA exits the nucleosome and is reminiscent to positions of TF binding sites located *in vivo* (43–45). As Gal4-VP16 is titrated with these Cy3-Cy5 labeled nucleosomes, Gal4-VP16 binds to its site within the nucleosome via the nucleosome site exposure mechanism (**Figure 3B**). This mechanism involves the nucleosome partially unwrapping, which transiently exposes the Gal4-VP16 target site. Gal4-VP16 then can bind to the site trapping the nucleosome in a partially unwrapped state. This results in a significant reduction in FRET efficiency.

We carried out FRET efficiency measurements of Gal4-VP16 titrations. As expected, the FRET efficiency is reduced as Gal4-VP16 traps the nucleosome in a partially unwrapped state (**Figure 3C**). We fit the FRET efficiency measurements to a binding isotherm to determine the binding half saturation concentration, S_1/2_, which is interpreted as an apparent K_D_. We find that Gal4-VP16 binds to the target sequence within the nucleosome with an S_1/2_ of 55 ± 2 nM (**Figure 3D**). To control for the impact of the VP-16 AD we also carried out titrations with Gal4-DBD alone and determined an S_1/2_ of 19 ± 1 nM. This shows that Gal4-VP16 is able to bind to its target site within nucleosomes. This binding appears to follow the site exposure model (4) even with the additional steric bulk of the VP16 activator, although with a 3-fold higher apparent K_D_.

### The transcription activator Gal4-VP16 significantly enhances the acetylation of nucleosomes by the HAT modules of ATAC and SAGA

Having determined that (i) the ATAC_HAT_ and SAGA_HAT_ interact directly with nucleosomes, (ii) the VP16 activator domain directly interacts with ATAC_HAT_ and SAGA_HAT_, and (iii) Gal4-VP16 binds efficiently within nucleosomes via the site exposure model, we decided to investigate the influence of Gal4-VP16 on the ability for ATAC_HAT_ and SAGA_HAT_ to acetylate nucleosomes within histone H3 and H4 tails. Nucleosomes were incubated for 30 minutes with an increasing concentration of either ATAC_HAT_ or SAGA_HAT_, Ac-CoA, and either with or without Gal4-VP16. The HAT module concentrations were increased over the concentration at which they bind nucleosomes (**Figure 1**). A Gal4-VP16 concentration of 160 nM was used to ensure saturated binding since it is approximately 2.5-fold higher than the S_1/2_ measured by the above FRET measurements (**Figure 3D**). Western blot assays were used to quantify the acetylation of the H3 tail at lysine 9 (H3K9) and the H4 tail at lysine 5 (H4K5) with and without Gal4-VP16 (**Figure 4**). To control for nucleosome loading we relied on Cy5 fluorescence, which is attached to H2A at K119C.

**Figure 4:**
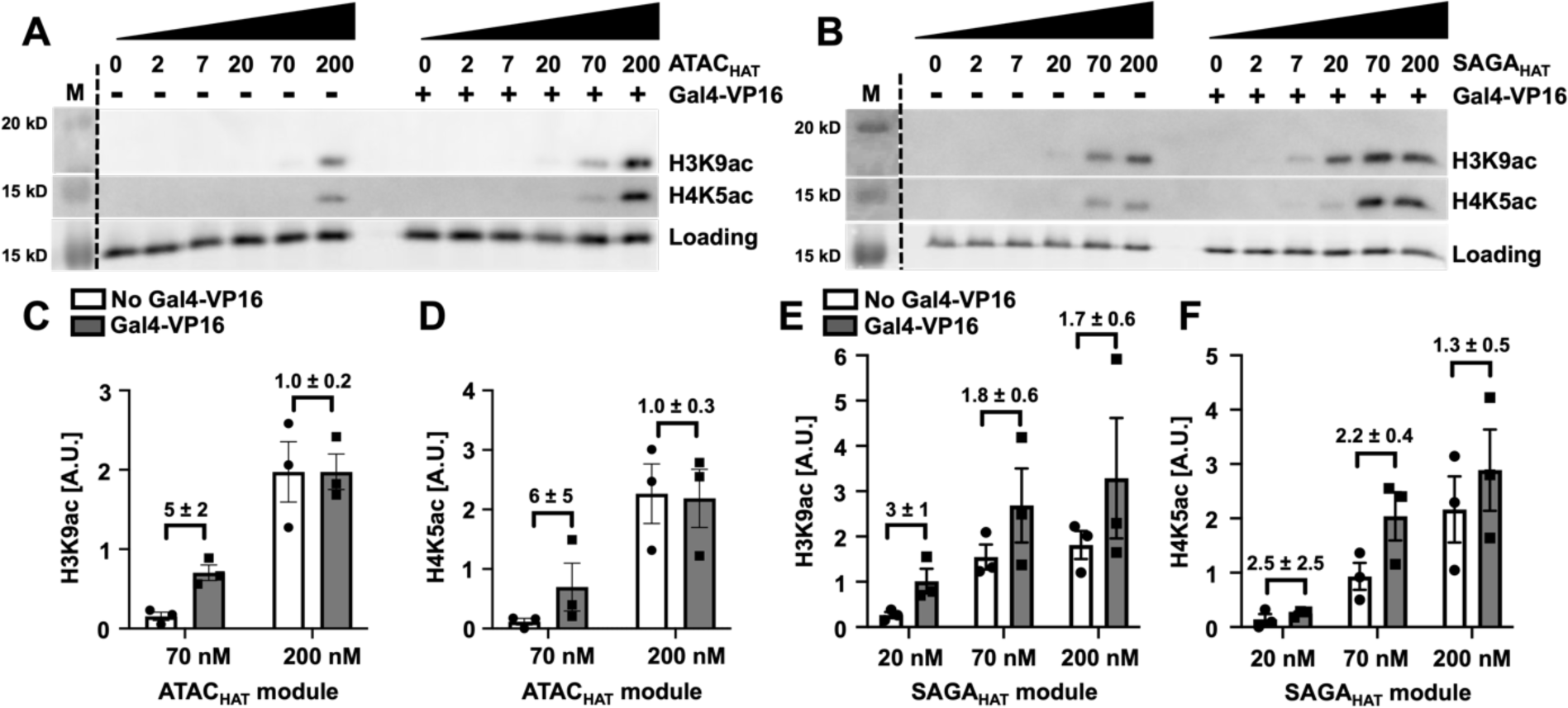
Gal4-VP16 influences the acetylation of the SAGA HAT module. Acetylation-coupled western blot assay of the ATAC_HAT_ (A) and SAGA_HAT_ (B) module at increasing concentrations with and without Gal4-VP16 at the indicated PTM sites. The fluorescence of Cy5-labeled histone H2A within the nucleosome was used to quantify nucleosome loading. Acetylation efficiency was tested by western blot using antibodies recognizing either histone H3K9ac, or H4K5ac. Images are representative of three independent experiments with similar results (*n* = 3). Two-tailed, unpaired t-test with Welch’s correction was conducted using Prism 9 software. Error bars represent standard error of the mean (SEM). (C,D) Bar plot of A. (E,F) Bar plot of B.

We find that Gal4-VP16 enhances acetylation of both H3K9 and H4K5 by ATAC_HAT_ (**Figure 4A-C**). At sub-saturating concentrations of 70 nM ATAC_HAT_, Gal4-VP16 induced a 5-fold and 6-fold increase in the acetylation of H3K9 and a H4K5, respectively (**Figure 4B-C, Supplementary Table S2**). In contrast, the impact of Gal4-VP16 on acetylation by SAGA_HAT_ was less dramatic. At the sub-saturating concentration of 20 nM SAGA_HAT_, Gal4-VP16 increased acetylation activity at H3K9 by 3-fold (**Figure 4E**), while at the H4K5 location, Gal4-VP16 increased acetylation activity modestly. At 20 nM of SAGA_HAT_, Gal4-VP16 did not induce a statistically significant increase in acetylation of H4K5, while a statistically significant change in acetylation of H4K5 (2.2-fold) with 70 nM SAGA_HAT_ and Gal4-VP16 was observed.

These observed differences in acetylation by ATAC_HAT_ and SAGA_HAT_ are consistent with *in vivo* studies, where SAGA targets H3 at K9 and K14 (17, 21, 22), while ATAC appears to acetylate both histone H3 and H4 (16, 23–27). In addition, the difference in the impact of Gal4-VP16 on acetylation by ATAC_HAT_ could in part be due to the lower starting activity of ATAC_HAT_ as compared to SAGA_HAT_, allowing Gal4-VP16 to have a larger impact on ATAC_HAT_ acetylation activity. Overall, these results indicate that Gal4-VP16, which directly interacts with both HAT modules and nucleosomes that contain the Gal4 binding site, facilitates acetylation of both the H3 and H4 tails by both HAT modules, although Gal4-VP16 has a larger impact on ATAC_HAT_ targeted acetylation.

### Trapping the nucleosome in a partially unwrapped state enhances the ATAC HAT module acetylation activity but suppresses the SAGA HAT module acetylation activity

The increase in HAT module acetylation activity by Gal4-VP16 could occur by two non-mutually exclusive mechanisms. The first mechanism is that the direct interaction of the VP16 AD helps stabilize the HAT module binding to the nucleosome, which in turn enhances the acetylation activity. The second mechanism is that Gal4-VP16 traps the nucleosome in a partially unwrapped state, increasing the exposure of the H3 and H4 tails for HAT module binding and acetylation. To determine the impact of nucleosome unwrapping alone, we carried out the above-described acetylation activity measurements with Gal4-DBD alone, which does not contain the VP16 AD. Gal4-DBD reduces the FRET efficiency as it binds its target site within the nucleosome, demonstrating that it traps the nucleosome in a partially unwrapped state similarly to Gal4-VP16 (**Figure 3**). This allowed the determination of the impact of the DBD of a TF trapping the nucleosome in an unwrapped state on HAT module acetylation activity without the confounding impact of interactions with an activator domain.

As with Gal4-VP16, nucleosomes were incubated for 30 minutes with an increasing concentration of either ATAC_HAT_ or SAGA_HAT_, with Ac-CoA, and either with or without 70 nM Gal4-DBD, which ensures saturating binding of nucleosomes by Gal4-DBD (**Figure 3**). Western blot assays were again used to quantify the acetylation at H3K9 and H4K5. We observe minimal acetylation activity at H3K9 below 70 nM of ATAC_HAT_ and 20 nM SAGA_HAT_ (**Supplementary Figure S6**). Interestingly at 70 nM of ATAC_HAT_, which is near the apparent K_D_, the acetylation activity of H3K9 is increased 4-fold by Gal4-DBD. At the saturating concentration of 200 nM ATAC_HAT_, Gal4-DBD no longer influences the acetylation activity at H3K9 (**Supplementary Figure S6**, **Supplementary Table S3**). In contrast, we find that Gal4-DBD does not induce a statistically significant change in the acetylation activity of ATAC_HAT_ at H4K5. For SAGA_HAT_, we found that Gal4-DBD did not enhance the acetylation activity of SAGA_HAT_ at either H3K9 or H4K5 (**Supplementary Figure 6, Supplementary Table S3**). Instead, the acetylation activity was reduced by about a factor of 2 at H3K9 with 70 nM SAGA_HAT_, while the change in acetylation activity at H4K5 was not statistically significant at any SAGA_HAT_ concentration.

These results are consistent with the structure of the nucleosome. The H3 tail extends out from the nucleosome where the DNA exits the nucleosome, which positions the H3 tail such that a partial nucleosome unwrapping could impact its accessibility. In contrast, the H4 tails extends out from the face of the histone octamer, which is at a position where partial nucleosome unwrapping is not likely to enhance H4 tail accessibility. Interestingly, these findings also suggest that trapping the nucleosome in a partially unwrapped state with Gal4-DBD provides ATAC_HAT_ additional access to the H3 tail to acetylate H3K9 but suppresses H3 tail access to SAGA_HAT_. This points to a possible mechanism where a TF can differentially target histone H3 tail acetylation by these two HAT modules.

### ATAC and SAGA HAT modules influence transcription factor accessibility in an acetylation-independent manner

Co-activator complexes are reported to be recruited to promoter regions by transcription factors to aid Pol II transcription initiation (68). However, the mechanisms by which coactivator complexes and transcription factors function together is still not fully understood. In the acetylation assays, we showed that the addition of Gal4-VP16 can significantly enhance the acetylation function of the studied HAT modules. This finding raised the question: do the HAT modules in turn influence the accessibility of Gal4-VP16 to the nucleosome? To investigate how ATAC_HAT_ and SAGA_HAT_ function with a TF to target nucleosomes, we used ensemble FRET efficiency measurements to quantify Gal4-VP16 binding within partially unwrapped nucleosomes in the presence or absence of either the ATAC_HAT_ or SAGA_HAT_ (**Figure 5A**, **D**). By comparing the Gal4-VP16 S_1/2_ for binding to nucleosomes with and without the HAT modules, we determined the impact of the HAT modules on the site accessibility for Gal4-VP16 occupancy at its site within a partially unwrapped nucleosome.

**Figure 5:**
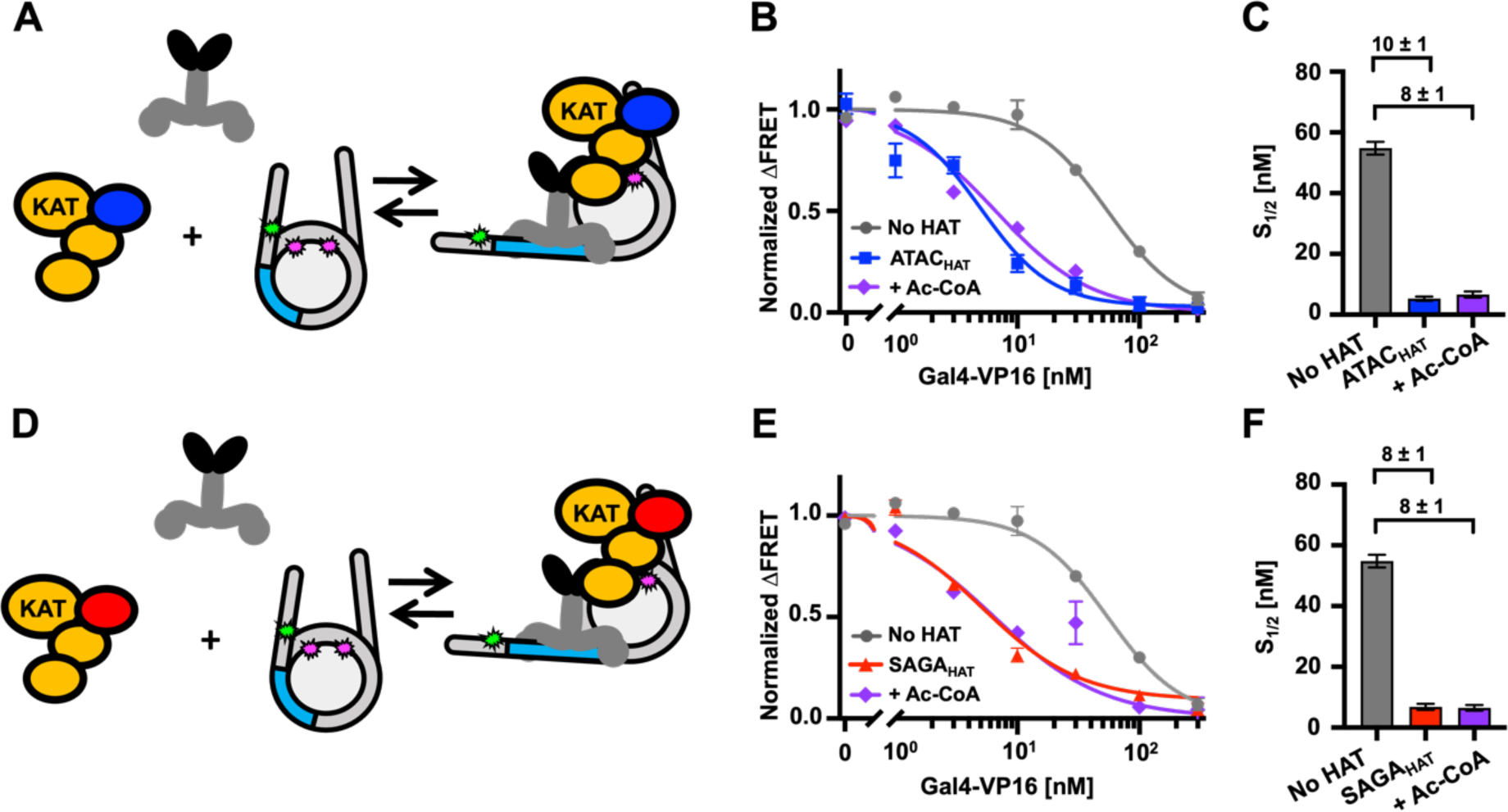
ATAC_HAT_ and SAGA_HAT_ influence nucleosome accessibility to Gal4-VP16 binding independent of Ac-CoA. (A) Model of Gal4-VP16 and the ATAC_HAT_ module binding to a nucleosome. (B) Normalized ΔFRET of Gal4-VP16 binding nucleosomes with increasing amounts. Gal4-VP16 binding to nucleosomes at a S_1/2_ of 55 ± 2 nM (dark gray circles), Gal4-VP16 binding to nucleosomes with the addition of 200 nM ATAC_HAT_ at a S_1/2_ of 5.3 ± 0.7 nM (dark blue squares), and with the addition of 30 uM Ac-CoA at a S_1/2_ of 7 ± 1 nM (purple diamond). (C) Bar plot of the S_1/2_ from B. (D) Model of Gal4-VP16 and the SAGA_HAT_ module binding to a nucleosome. (E) Normalized ΔFRET of Gal4-VP16 binding to nucleosomes at increasing amounts. Gal4-VP16 binding to nucleosomes at a S_1/2_ of 55 ± 2 nM (dark gray circles), Gal4-VP16 binding to nucleosomes with the addition of 150 nM SAGA_HAT_ at a S_1/2_ of 7 ± 1 nM (red triangles), and with the addition of 30 μM Ac-CoA at a S_1/2_ of 6.5 ± 0.9 nM (purple diamond). (F) Bar plot of S_1/2_ and standard error values from the weighted fits to binding isotherms in E.

We used HAT module concentrations of 200 nM and 150 nM for ATAC_HAT_ or SAGA_HAT_, respectively, to ensure that the HAT modules were at concentrations that efficiently bind nucleosomes (**Figure 1D**). We carried out these measurements without Ac-CoA to eliminate the impact of histone acetylation on Gal4-VP16 binding. In the presence of ATAC_HAT_, the binding affinity of Gal4-VP16 increased by 10-fold where the S_1/2_ lowered from 55 ± 2 nM to 5.3 ± 0.7 nM (**Figure 5B-C, Supplementary Table S4**). Likewise, the addition of SAGA_HAT_ increased the ability of Gal4-VP16 to bind by 8-fold where the S_1/2_ of Gal4-VP16 lowered to 7 ± 1 nM (**Figure 5E-F, Supplementary Table S4**). We then repeated these measurements with 30 μM Acetyl-CoA to determine how histone acetylation influences the impact of the HAT modules on Gal4-VP16 binding. We found that the addition of Ac-CoA did not statistically impact in the binding affinity of Gal4-VP16 (**Figure 5C**, **F**), which implies that the order of magnitude increases in Gal4-VP16 binding within the nucleosome by ATAC_HAT_ and SAGA_HAT_, but this increase is acetylation independent.

Since VP16 AD directly interacts with both HAT modules, we investigated if the VP16 AD is required for the acetylation independent increase in Gal4-VP16 binding by determining if the HAT modules facilitate Gal4-DBD binding and by how much. We carried out Gal4-DBD titrations with and without ATAC_HAT_ or SAGA_HAT_ to determine the change in the S_1/2_ of Gal4-DBD binding (**Supplementary Figure S7**, **Supplementary Table S4**). We found that ATAC_HAT_ increased Gal4-DBD binding by about 2.5-fold, where the S_1/2_ decreases from 19 ± 1 nM to 7.6 ± 0.8 nM, while SAGA_HAT_ increases Gal4-DBD binding by about 4-fold where the S_1/2_ decreased to 4.9 ± 0.3 nM. We also investigated the impact of Ac-CoA and found that this increase in Gal4-DBD binding was not significantly altered by the acetylation activity of either HAT module, implying that the enhanced binding of Gal4-DBD by both HAT modules is acetylation independent.

In combination, these results indicate that the VP16 AD is important for the order of magnitude increase of Gal4-VP16 binding induced by either ATAC_HAT_ or SAGA_HAT_. This is likely due to direct interactions between VP16 and the HAT modules. However, the more modest increase binding of Gal4 DBD alone indicates that while the HAT modules do not directly induce nucleosome unwrapping (**Figure 1E**, **Supplementary Figure S5B**), they can help stabilize Gal4 binding even without directly interacting with an activator domain, perhaps by preventing rewrapping. Overall, these findings indicate that not only do transcription activating TFs, such as Gal4-VP16, enhance ATAC_HAT_ and SAGA_HAT_ acetylation, but these HAT modules also facilitate TF binding independent of the acetylation activity.

## DISCUSSION

This study provides quantitative and mechanistic insight into how the HAT modules of two important co-activators, ATAC and SAGA, function with a transcription activating TF to target and open chromatin. Our results reveal that both ATAC_HAT_ and SAGA_HAT_ have two distinct yet synergistic functions: (i) coordinate with TFs to acetylate histones to open chromatin and (ii) target the same TFs to its binding site within the nucleosome. These observations also provide insight into how both the ATAC and SAGA full complexes function with a TF. KAT2A and other SAGA subunits have been reported to interact with transcription activating TFs (36), while ATAC subunit(s) that directly interacts with TFs has yet to be characterized. Interestingly, our *in vitro* binding experiments suggest that KAT2A in both ATAC_HAT_ and SAGA_HAT_ is likely to contribute to TF binding, and consequently to the recruitment of these HAT modules, and/or their holo complexes, to activate genes. In addition, the SAGA holo complex contains the TRRAP subunit that also interacts with TFs (38). In a recent screen test of 109 human TFs’ interactions with other proteins by proximity-dependent biotinylation (BioID) and affinity purification mass spectrometry (AP-MS) measurements, the authors described that the endogenous ATAC complex interacts with E2F1; ELF1, HNF4a, KLF3, KLF12, MYC; PPARγ, TYY1 TFs, while endogenous SAGA complex was shown to interact with KLF6, MYC, and MYOD TFs (69). Our results indicate that the ATAC and SAGA HAT modules may contribute to these TF interactions, which could help target the ATAC and/or SAGA holo complexes to promoters and enhancers. In addition, these HAT modules could also function outside of the endogenous ATAC and SAGA holo complexes (36, 37). Thus, it is conceivable that TFs can directly target the HAT modules, free or incorporated in their respective holo complexes, to TF target sites allowing the ATAC_HAT_ and SAGA_HAT_ modules or their holo complexes to function as transcription co-activators.

Our studies of the impact of Gal4-VP16 vs. Gal4-DBD on the acetylation activities of ATAC_HAT_ and SAGA_HAT_ allowed us to decouple the impact of the VP16 activator direct interaction with the HAT modules and the impact of the TF trapping the nucleosome in a partially unwrapped state. The combined results of Gal4-VP16 and Gal4-DBD indicate that a TF with an activator domain that directly interacts with the HAT module enhances ATAC_HAT_ acetylation of the H3 tail by 2 mechanisms: (i) it facilitates recruitment of ATAC_HAT_ module through direct interactions with the activator domain, and (ii) it increases ATAC_HAT_ accessibility to the histone tail through partial nucleosome unwrapping. In contrast, only the first mechanism appears to be important for ATAC_HAT_ acetylation of the H4 tail, which as mentioned above is consistent with the location of the H4 tail within the nucleosome. For SAGA_HAT_, trapping the nucleosome in a partially unwrapped state does not impact its acetylation activity, implying that only the first mechanism has a significant impact on SAGA_HAT_ acetylation activity of both the H3 and H4 tails. These results highlight differences between the two HAT modules, which could be important in how ATAC_HAT_ and SAGA_HAT_ target distinct chromatin locations.

The new function of ATAC_HAT_ and SAGA_HAT_ modules identified in this study by *in vitro* experiments, where they help target a TF to its recognition site within a nucleosome, is highly synergistic with their well-established function to acetylate histones. By helping the TF target their binding sites within nucleosomes, ATAC_HAT_ and SAGA_HAT_ enhance the targeting for their own HAT activity. This targeting could happen through three non-exclusive models (**Figure 6**) where either the TF and nucleosome bind first (Model 1), the TF and the HAT module bind first (Model 2), or the HAT module and the nucleosome bind first (Model 3). Since we found that Gal4-VP16, HAT modules, and nucleosomes all interact strongly with each other, it is possible that all three pair wise interactions are possible pathways to the nucleosome being bound by both Gal4-VP16 and a HAT module. Our current studies do not differentiate between these pathways. However, a previous study reported that suppression of Gal4-DBD occupancy at its site within the nucleosome is dominated by nucleosome accelerated dissociation (70). This suggests that when the nucleosome is bound by Gal4-VP16 and the HAT module, both could have a slower dissociation rate, which would result in increased dissociation equilibrium constant irrespective of the dominate model(s) in **Figure 6**. Future studies are needed to investigate the key states that allow both HAT modules to increase Gal4 binding by an order of magnitude.

**Figure 6.**
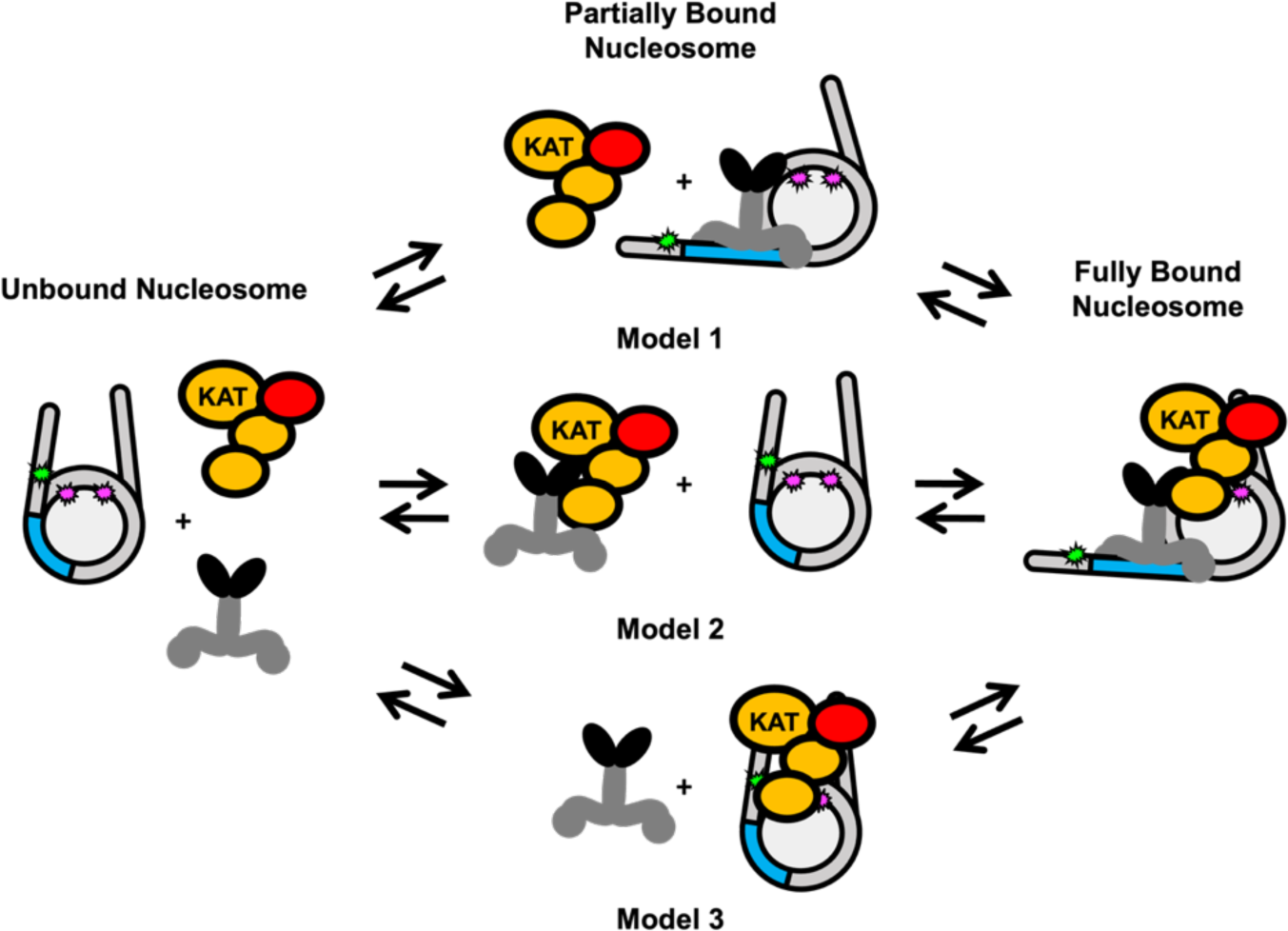
Models of HAT module and TF binding to a partially unwrapped nucleosome. Model 1: Gal4-VP16 first binds to its recognition sequence and then recruits the HAT module. Model 2: Gal4-VP16 and the HAT module interact with each other first before binding the nucleosome together. Model 3: the HAT module interacts with the nucleosome first, allowing Gal4-VP16 to bind at a higher affinity. Since the impact of the HAT module on TF binding is acetylation independent, acetylation is not indicated in the model. However, histone acetylation will occur when the HAT module is bound to the nucleosome, which includes the fully bound nucleosome and the intermediate state of Model 3.

Interestingly, recent molecular and genetic studies revealed that many histone-modifying enzymes have functions independent of their HAT, or methyltransferase activities. Many of the histone modifications deposited by these enzymes, previously considered to be crucial for transcriptional activation, have been demonstrated to function for gene expression independent of their catalytic activities (71–73). Our discovery of the new HAT-independent function of the ATAC and SAGA HAT modules in targeting TFs to their recognition sites within chromatin in a HAT-independent manner, suggests that the newly discovered function of these chromatin-modifying complexes may be even more important than their given enzymatic activities. Future experiments will be required to verify whether the HAT-independent targeting role of these complexes also operates in living organisms.

## ACKNOWLEDGEMENTS

We thank the members of the Poirier, Tora, and Berger labs for helpful discussions and comments. We are grateful the Parthun lab and the Musier-Forsyth labs for help with the Western Blots. This work was supported by the National Institutes of Health (R35 GM139564 to M.G.P. and L.T., and R01 GM131626), and the Agence Nationale de la Recherche (ANR) (ANR-19-CE11-0003-02 to L.T., ANR-20-CE12-0017-03 to L.T., ANR-PRCI-19-CE12-0029-01 to L.T., ANR-22-CE11-0013-01_ACT; to L.T.). As part of the ITI 2021-2028 program of the University of Strasbourg, this work was also supported by IdEx Unistra (ANR-10-IDEX-0002), and by SFRI-STRAT’US project (ANR 20-SFRI-0012) and EUR IMCBio (ANR-17-EURE-0023) under the framework of the French Investments for the Future Program. IB is supported by the Wellcome Trust (106115/Z/14/Z).

## DATA AVAILABILITY

The experimental data sets are either included in this manuscript, the supplemental information, or are available from the authors upon request.

## CONTRIBUTIONS

K.Chesnutt, M.G.P., and L.T. conceived and designed the research. K.Chesnutt carried out the FA, Western blot, and FRET experiments. K.Chesnutt and K.Cox prepared the nucleosome samples. G.Y. carried out the pull-down experiments. C.T. and G.G. prepared the ATAC and SAGA HAT modules. M.G.P. and L.T. supervised the study. K.Chesnutt, M.G.P. and L.T. wrote the first draft. All authors edited and finalized the manuscript.

## Supplemental Figures

**Figure S1.**
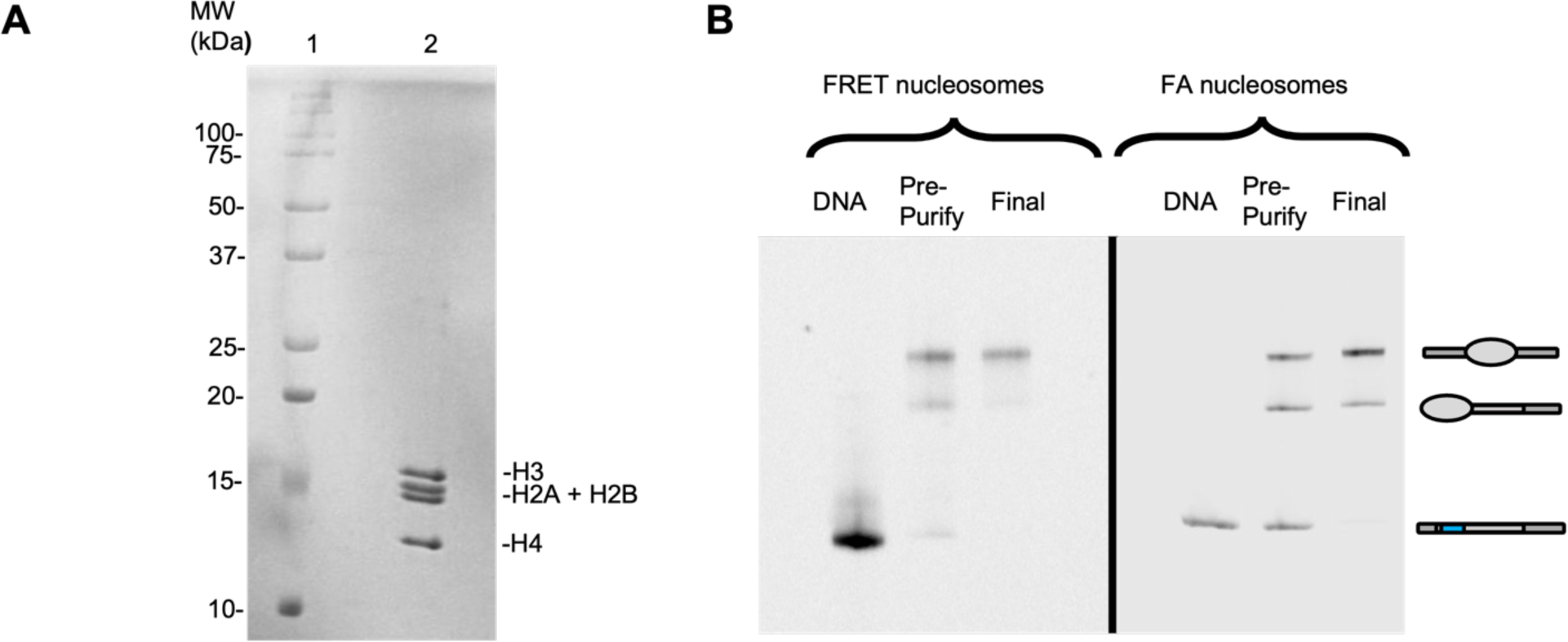
Histone octamer and nucleosome preparations. (A) Purified histone octamer visualized on SDS PAGE gel (lane 2). Lane 1: molecular weight (MW) markers in kDa. (B) Native PAGE analysis of purified nucleosomes. DNA lane contains only DNA used to make nucleosomes. Pre-Purify lane contains nucleosomes before sucrose gradient purification. Final lane contains nucleosomes after sucrose gradient purification. The slower mobility nucleosome band contains center-positioned nucleosomes while the faster mobility nucleosome band contains end-positioned nucleosomes.

**Figure S2.**
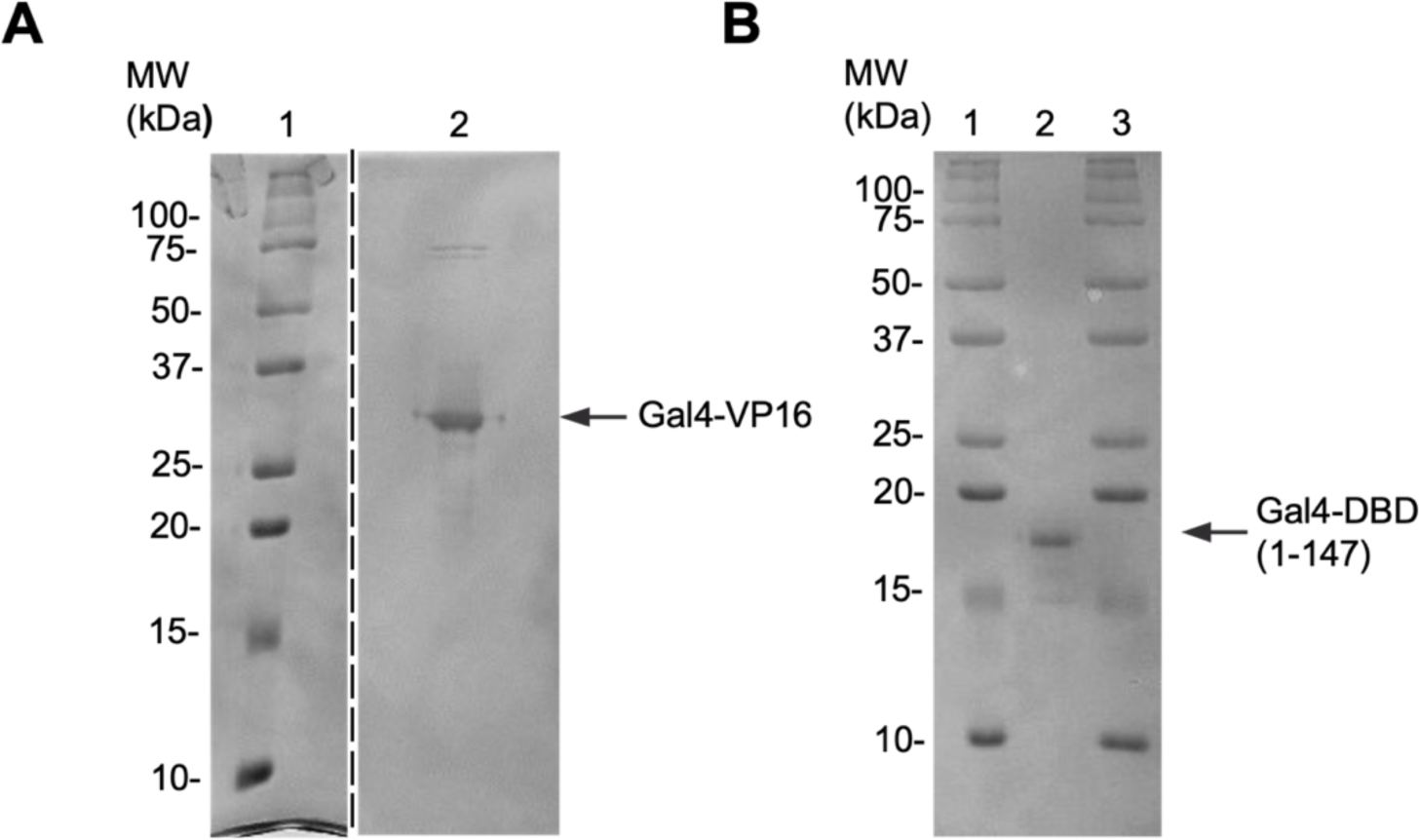
Gal4-VP16 and Gal4-DBD purification. The purified Gal4-VP16 (A) and Gal4-DBD (from amino acid 1 to 147) (B) proteins were visualized on SDS PAGE gel by Coomassie blue staining, Lane 1 in (A) and lanes 1 and 3 in (B): molecular weight (MW) markers in kDa.

**Figure S3.**
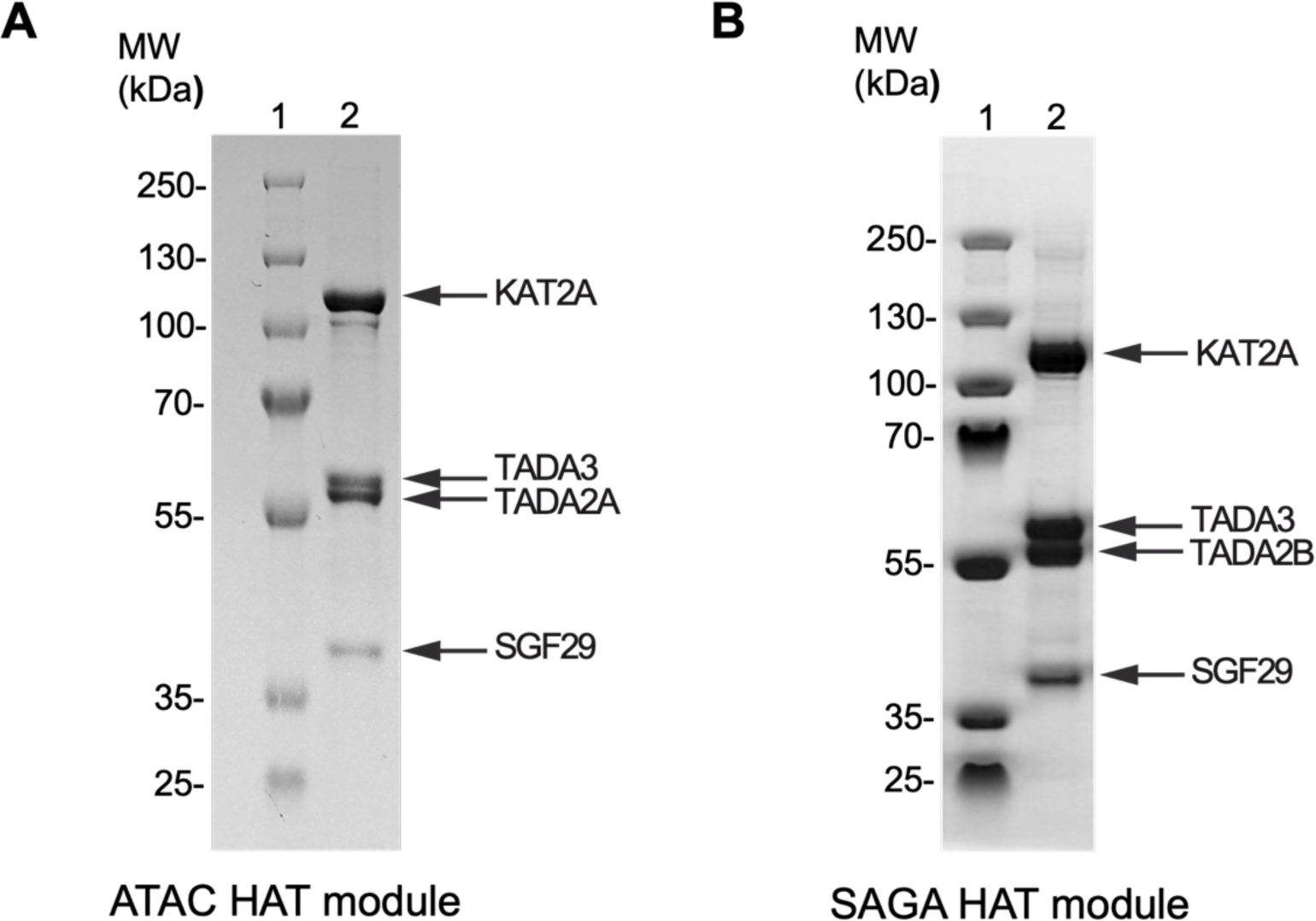
ATAC and SAGA HAT module purifications. (A) Highly purified ATAC_HAT_ (A) and SAGA_HAT_ (B) were visualized on SDS PAGE gel by Coomassie blue staining. Lane 1 in (A) and (B) :molecular weight (MW) markers in kDa.

**Figure S4.**
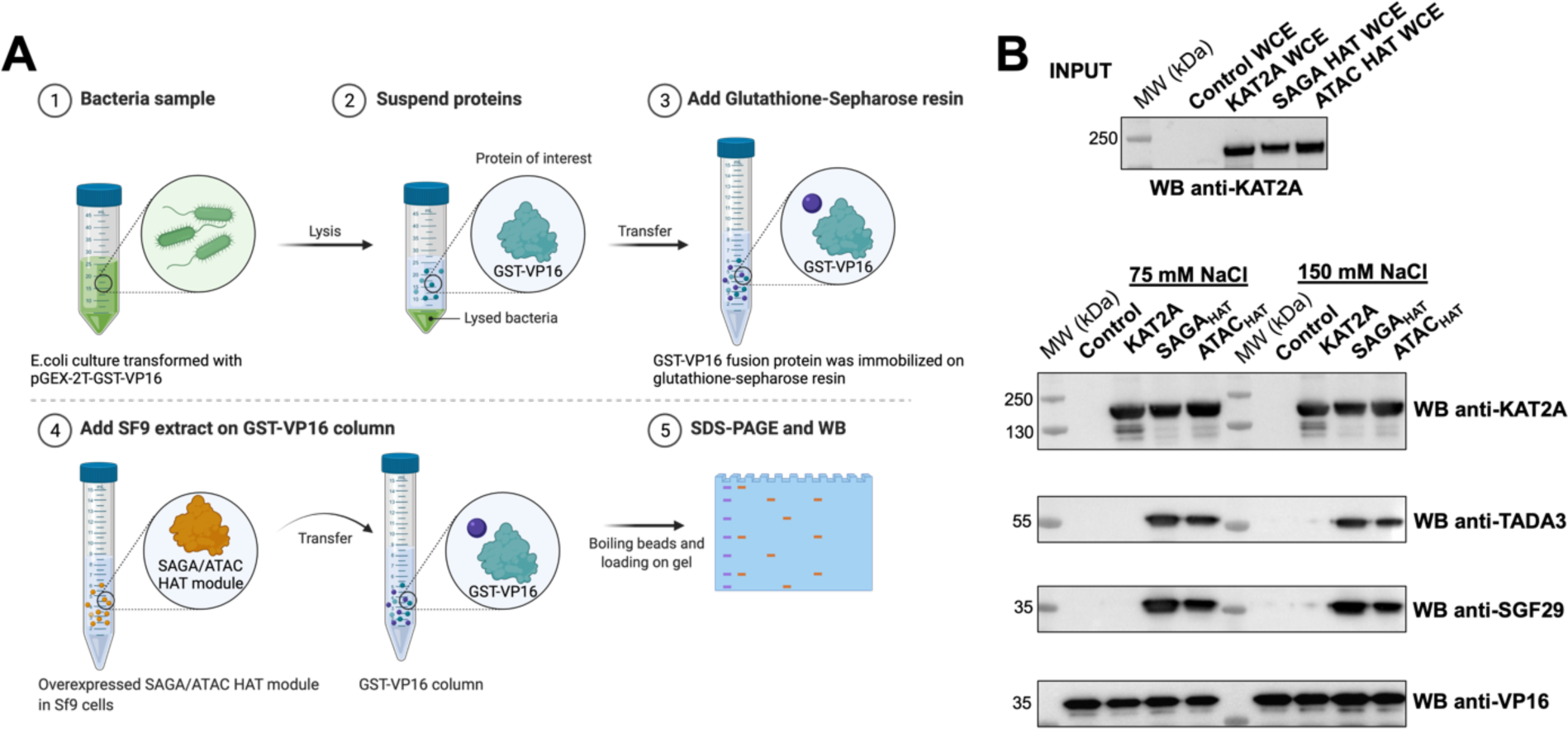
Specific binding of the HAT module of SAGA and ATAC complexes to VP16-AD. (A) Schematic representation of experimental workflow. (B) Western blot analysis of the elutes from the GST-VP16 column, using SF9 whole cell extracts (WCE) where recombinant KAT2A protein, SAGA_HAT_ or ATAC_HAT_ modules were overexpressed by baculovirus overexpression system in Sf9 cells, separately. Upper panel indicates overexpression levels of KAT2A in each whole cell extracts. Lower panels indicate the presence the HAT module subunits in elutions from the GST-VP16 column in the presence of either 75 mM NaCl or 150 mM NaCl. Molecular weight (MW) markers in kDa are indicated. The indicated proteins were detected by Western blot assays (WB) using antibodies raised against the indicated proteins.

**Figure S5.**
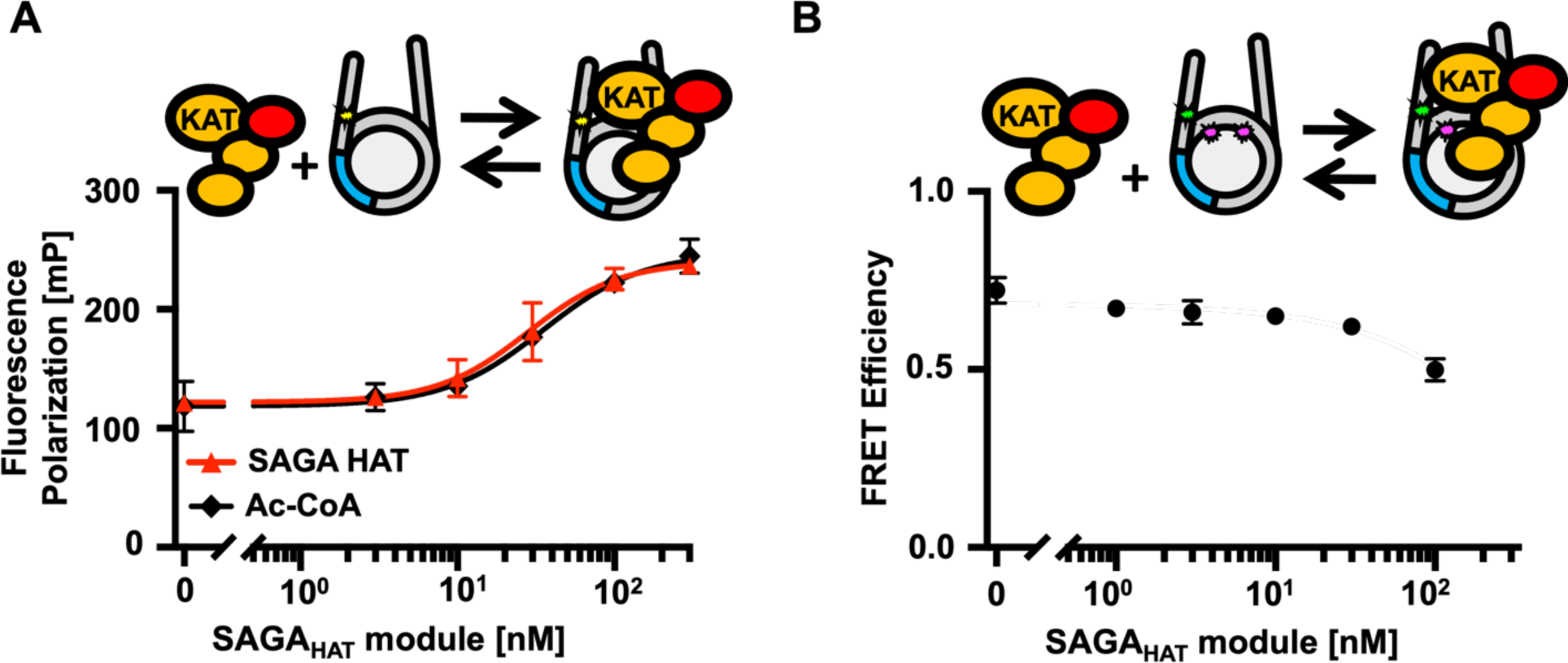
SAGA_HAT_ interactions with nucleosomes. (A) Fluorescence Polarization in mP of nucleosomes with increasing amounts of the SAGA_HAT_ module. The SAGA_HAT_ module binds with an S1/2 of 25.0 ± 0.3 nM (red triangle) and reaches saturating conditions around 100 nM. The SAGA_HAT_ module with Ac-CoA binds at a S1/2 of 37 ± 3 nM. (B) FRET of the SAGA_HAT_ module binding nucleosomes at increasing amounts. n = 3

**Figure S6:**
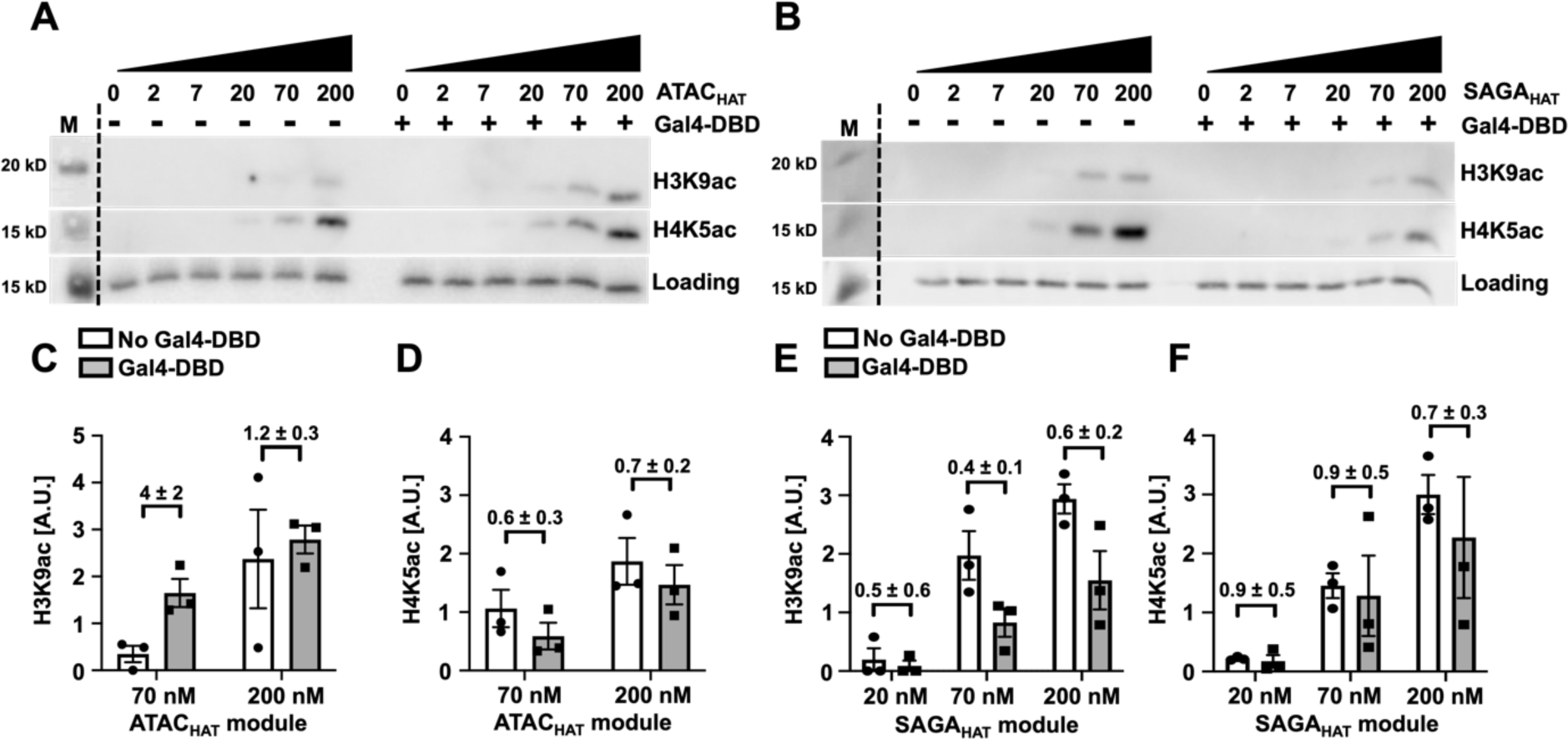
Influence of the of Gal4-DBD on the acetylation by the ATAC and SAGA HAT modules. Acetylation assay by the ATAC_HAT_ (A) or SAGA_HAT_ (B) modules at increasing concentrations with and without Gal4-DBD at the indicated PTM sites. The fluorescence of Cy5-labeled histone H2A within the nucleosome was used to observe nucleosome loading. Acetylation efficiency was tested by Western blot using antibodies recognizing either histone H3K9ac or H4K5ac. Images are representative of three independent experiments with similar results (*n* = 3). Two-tailed, unpaired t-test with Welch’s correction was conducted using Prism 9 software. Error bars represent standard error of the mean. (C, D) Bar plots of A (E, F) Bar plots of B.

**Figure S7:**
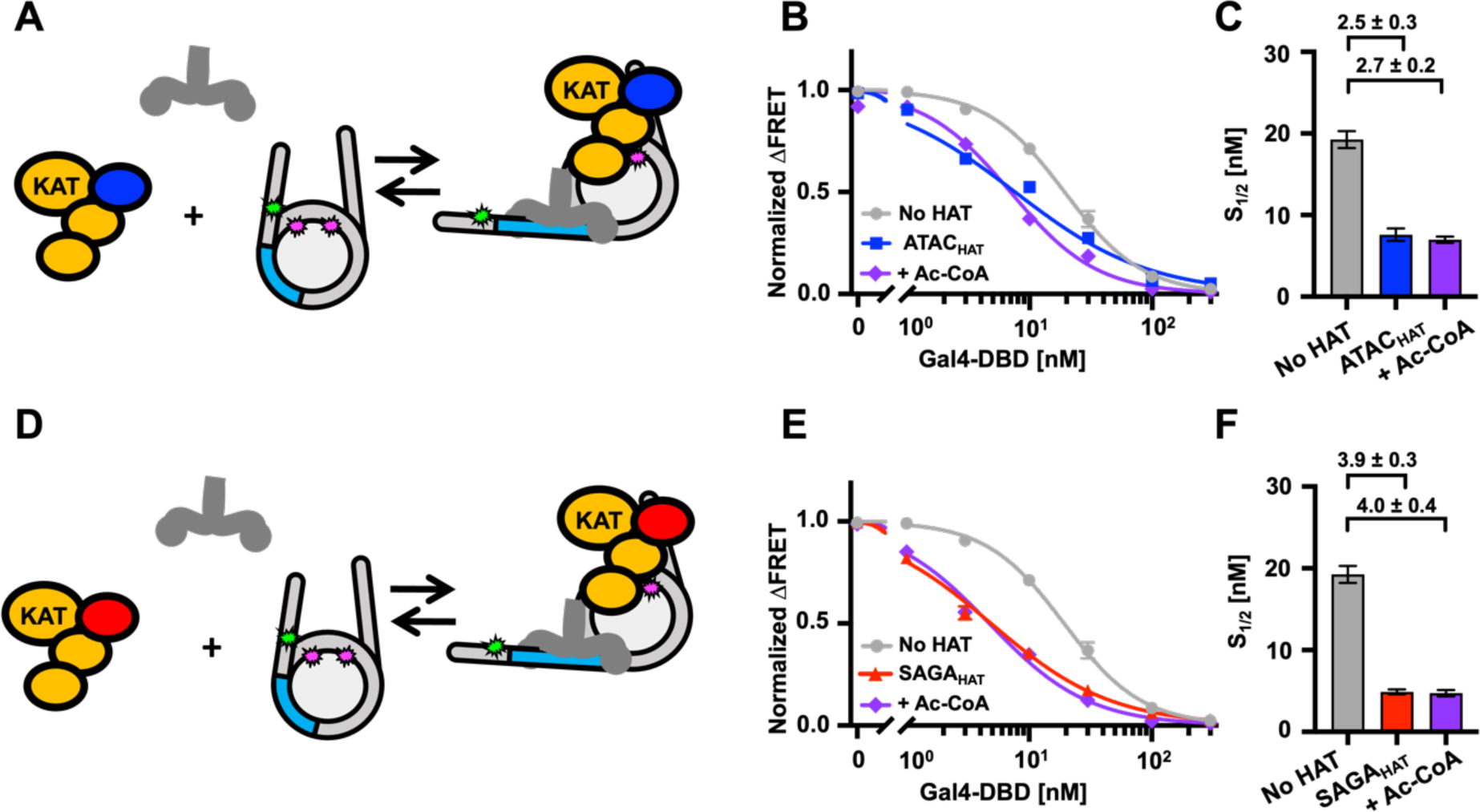
ATAC_HAT_ and SAGA_HAT_ influence nucleosome accessibility Gal4-DBD binding independent of Ac-CoA. (A) Binding model of Gal4-DBD and the ATAC_HAT_ module binding to a nucleosome. (B) Normalized ΔFRET of Gal4-DBD binding nucleosomes with increasing amounts. Gal4-DBD binding to nucleosomes at a S_1/2_ of 19 ± 1 nM (light gray circles), Gal4-DBD binding to nucleosomes with the addition of 200 nM ATAC_HAT_ at a S_1/2_ of 7.6 ± 0.8 nM (dark blue squares), and with the addition of 30 µM Ac-CoA at a S_1/2_ of 7.0 ± 0.4 nM (purple diamond). (C) Bar plot of S1/2 values from B (D) Binding model of Gal4-DBD and the SAGA_HAT_ module binding to a nucleosome. (E) Normalized ΔFRET of Gal4-DBD binding to nucleosomes at increasing amounts. Gal4-DBD binding to nucleosomes at a S_1/2_ of 19 ± 2 nM (light gray circles), Gal4-VP16 binding to nucleosomes with the addition of 150 nM SAGA_HAT_ at a S_1/2_ of 4.9 ± 0.3 nM (red triangles), and with the addition of 30 μM Ac-CoA at a S_1/2_ of 4.7 ± 0.4 nM (purple diamond). (F) Bar plot of S_1/2_ and standard error values from the weighted fits to binding isotherms in E.

## Supplemental Tables

**Table S1.** Fluorescence polarization fit values. Error indicate +/- 1 standard error.

| HAT Module | Ac-CoA | S<sub>1/2</sub> | P value | P < 0.2 |
| --- | --- | --- | --- | --- |
| ATAC HAT | No | 23 ± 4 | 0.49 | n.s. |
| ATAC HAT | Yes | 19.6 ± 0.6 |  |  |
| SAGA HAT | No | 25.0 ± 0.3 | 0.06 | yes |
| SAGA HAT | Yes | 37 ± 3 |  |  |

**Table S2.** Summary of ATAC_HAT_ and SAGA_HAT_ Western blot analysis with Gal4-VP16. Error indicate +/- 1 standard error.

| HAT (nM) | TF | PTM | Blot Signal | Ratio | P value | P < 0.2 |
| --- | --- | --- | --- | --- | --- | --- |
| ATAC (70 nM) | + Gal4-VP16 | H3K9ac | 0.7 ± 0.1 | 5 ± 2 | 0.01 | yes |
|  | - Gal4-VP16 | H3K9ac | 0.15 ± 0.05 |  |  |  |
| ATAC (200 nM) | + Gal4-VP16 | H3K9ac | 2.0 ± 0.2 | 1.0 ± 0.2 | 1.00 | n.s. |
|  | - Gal4-VP16 | H3K9ac | 2.0 ± 0.3 |  |  |  |
| ATAC (70 nM) | + Gal4-VP16 | H4K5ac | 0.7 ± 0.4 | 6 ± 5 | 0.28 | n.s. |
|  | - Gal4-VP16 | H4K5ac | 0.11 ± .06 |  |  |  |
| ATAC (200 nM) | + Gal4-VP16 | H4K5ac | 2.2 ± 0.5 | 1.0 ± 0.3 | 0.91 | n.s. |
|  | - Gal4-VP16 | H4K5ac | 2.3 ± 0.5 |  |  |  |
| SAGA (20 nM) | + Gal4-VP16 | H3K9ac | 1.0 ± 0.3 | 3 ± 1 | 0.10 | yes |
|  | - Gal4-VP16 | H3K9ac | 0.3 ± 0.1 |  |  |  |
| SAGA (70 nM) | + Gal4-VP16 | H3K9ac | 2.7 ± 0.8 | 1.8 ± 0.6 | 0.30 | n.s. |
|  | - Gal4-VP16 | H3K9ac | 1.5 ± 0.3 |  |  |  |
| SAGA (200 nM) | + Gal4-VP16 | H3K9ac | 3 ± 1 | 1.7 ± 0.6 | 0.38 | n.s. |
|  | - Gal4-VP16 | H3K9ac | 1.8 ± 0.3 |  |  |  |
| SAGA (20 nM) | + Gal4-VP16 | H4K5ac | 0.25 ± 0.03 | 2.5 ± 2.5 | 0.37 | n.s. |
|  | - Gal4-VP16 | H4K5ac | 0.1 ± 0.1 |  |  |  |
| SAGA (70 nM) | + Gal4-VP16 | H4K5ac | 2.0 ± 0.3 | 2.2 ± 0.4 | 0.11 | yes |
|  | - Gal4-VP16 | H4K5ac | 0.9 ± 0.08 |  |  |  |
| SAGA (200 nM) | + Gal4-VP16 | H4K5ac | 2.9 ± 0.8 | 1.3 ± 0.5 | 0.49 | n.s. |
|  | - Gal4-VP16 | H4K5ac | 2.2 ± 0.6 |  |  |  |

**Table S3.** Summary of ATAC_HAT_ and SAGA_HAT_ Western blot analysis with Gal4-DBD. Error indicate +/- 1 standard error.

| HAT (nM) | TF | PTM | Blot Signal | Ratio | P value | P < 0.2 |
| --- | --- | --- | --- | --- | --- | --- |
| ATAC (70 nM) | + Gal4-DBD | H3K9ac | 1.7 ± 0.3 | 4 ± 2 | 0.03 | yes |
|  | - Gal4-DBD | H3K9ac | 0.4 ± 0.2 |  |  |  |
| ATAC (200 nM) | + Gal4-DBD | H3K9ac | 2.8 ± 0.3 | 1.2 ± 0.3 | 0.74 | n.s. |
|  | - Gal4-DBD | H3K9ac | 2.4 ± 0.6 |  |  |  |
| ATAC (70 nM) | + Gal4-DBD | H4K5ac | 0.6 ± 0.3 | 0.6 ± 0.3 | 0.30 | n.s. |
|  | - Gal4-DBD | H4K5ac | 1.06 ± .08 |  |  |  |
| ATAC (200 nM) | + Gal4-DBD | H4K5ac | 1.9 ± 0.6 | 0.7 ± 0.2 | 0.49 | n.s. |
|  | - Gal4-DBD | H4K5ac | 2.8 ± 0.3 |  |  |  |
| SAGA (20 nM) | + Gal4-DBD | H3K9ac | 0.09 ± 0.09 | 0.5 ± 0.6 | 0.66 | n.s. |
|  | - Gal4-DBD | H3K9ac | 0.2 ± 0.2 |  |  |  |
| SAGA (70 nM) | + Gal4-DBD | H3K9ac | 0.8 ± 0.2 | 0.4 ± 0.1 | 0.09 | yes |
|  | - Gal4-DBD | H3K9ac | 2.0 ± 0.4 |  |  |  |
| SAGA (200 nM) | + Gal4-DBD | H3K9ac | 1.6 ± 0.5 | 0.6 ± 0.2 | 0.09 | yes |
|  | - Gal4-DBD | H3K9ac | 2.9 ± 0.3 |  |  |  |
| SAGA (20 nM) | + Gal4-DBD | H4K5ac | 0.2 ± 0.1 | 0.9 ± 0.5 | 0.70 | n.s. |
|  | - Gal4-DBD | H4K5ac | 0.22 ± 0.02 |  |  |  |
| SAGA (70 nM) | + Gal4-DBD | H4K5ac | 1.3 ± 0.7 | 0.9 ± 0.5 | 0.83 | n.s. |
|  | - Gal4-DBD | H4K5ac | 1.5 ± 0.2 |  |  |  |
| SAGA (200 nM) | + Gal4-DBD | H4K5ac | 2 ± 1 | 0.7 ± 0.3 | 0.56 | n.s. |
|  | - Gal4-DBD | H4K5ac | 3.0 ± 0.3 |  |  |  |

**Table S4.** Measurement values of transcription factor S_1/2_ with HAT modules. Error indicate +/- 1 standard error.

| TF | HAT Module | Ac-CoA | S<sub>1/2</sub> | S<sub>1/2</sub> noHAT / S<sub>1/2</sub> HAT | P value |
| --- | --- | --- | --- | --- | --- |
| Gal4-VP16 | None | - | 55 ± 2 | N.A. | N.A. |
| Gal4-VP16 | ATAC | - | 5.3 ± 0.7 | 10 ± 1 | 0.0006 |
| Gal4-VP16 | ATAC | + | 7 ± 1 | 8 ± 1 | 0.0003 |
| Gal4-VP16 | SAGA | - | 7 ± 1 | 8 ± 1 | 0.0003 |
| Gal4-VP16 | SAGA | + | 6.5 ± 0.9 | 8 ± 1 | 0.0003 |
| Gal4-DBD | None | - | 19 ± 1 | N.A. | N.A. |
| Gal4-DBD | ATAC | - | 7.6 ± 0.8 | 2.5 ± 0.3 | 0.001 |
| Gal4-DBD | ATAC | + | 7.0 ± 0.4 | 2.7 ± 0.2 | 0.003 |
| Gal4-DBD | SAGA | - | 4.9 ± 0.3 | 3.9 ± 0.3 | 0.003 |
| Gal4-DBD | SAGA | + | 4.7 ± 0.4 | 4.0 ± 0.4 | 0.002 |

